# House of Clocks: On the Misuse of Ageing Composite Measures

**DOI:** 10.1101/2025.05.24.655934

**Authors:** Ignophi Hu

## Abstract

**Objective:** This article examines the structural implications of composite variables developed in the field of “ageing”. First, using data from the Baltimore Longitudinal Study of Aging (BLSA), we illustrate how the property of feature convergence arises in practice. Second, we show how this constrains both cross-sectional stratification and longitudinal monitoring. Third, we extend the analysis to molecular implementations and demonstrate that similar structural issues persist. Fourth, we examine how these properties impact common objectives - intervention evaluation, prediction, and investigation - and show that the limitations are not confined to linear models. Finally, we consider additional properties that may help explain the widespread appeal and persistence of these models, despite clear conflicts with their stated objectives.

## 1. Introduction

A composite variable reduces sets of measurements into a single value by summing, averaging, weighting, or applying more sophisticated transformations [1]. These derived measures serve as simple models that offer a reduced representation of a multi-dimensional set. They are often used when the construct of interest is latent -that is, not directly observable - or when the underlying feature space is large, noisy, or difficult to interpret in its raw form. Composite variables are widely used in various fields. A prominent example is the Intelligence Quotient (IQ) [1, 2], a composite variable that aggregates a battery of cognitive test scores into a single metric intended to inform on the latent construct of “intelligence” or “cognitive ability”. Within the field of “ageing”, composite variables and their associated latent constructs have been especially popular. Over the decades, these have appeared under various labels, including functional age [3], biological age [4], healthspan metrics [5, 6], frailty indices [7], organ clocks [8], and others. The composite score and the latent construct are often referred to interchangeably by the same label [9] - for example, calling the empirical output simply “biological age” or “brain age”. While convenient, this practice blurs the distinction between the empirical score and the theoretical construct, complicating discussions of validity, interpretation, and utility.^**1**^

These composite variables have been built using features ranging from molecular to organismal [2, 10, 11, 12, 13] and span almost every type of statistical model - from simple summary scores to deep, multi-layer architectures [7, 14, 15]. Despite these differences, many implementations implicitly adopt the same theoretical generative model (**Figure 1**, **“Theoretical Model”**). This model posits a latent construct - commonly referred to as “functional state” or “biological age” - which, though unobservable, is assumed to give rise to a set of measurable outcomes. Some of these measures are more readily accessible. The implication of this model is that proximal measures can be used to estimate the latent construct, which in turn allows inference about distal outcomes - often the primary targets of interest [1].^**2**^ This explicit structural representation makes it possible to clarify several key assumptions embedded in the posited generative model - assumptions that directly influence its scientific and clinical utility:

- **(A1) Feature Convergence.** When a multidimensional set of input features is collapsed into a lower-dimensional (often scalar) composite, the mapping is necessarily many-to-one; consequently, multiple distinct biological configurations can yield identical composite scores. Although at the level of the theoretical model this appears as an assumption, in any statistical instantiation it is an unavoidable property rather than a matter of empirical choice.^**3**^
- **(A2) Sufficiency of the Composite** The latent construct is assumed to retain all (or nearly all) relevant information from the original features necessary for predicting distal outcomes. When the composite score is treated as a direct proxy for this latent construct, it is implicitly assumed to capture that same predictive sufficiency. As a result - and in conjunction with (A1) - it follows that individuals with the same composite value, despite having different input profiles, are expected to exhibit similar risks and clinical outcomes. Once the composite is known, the original features are assumed to contribute minimal additional predictive value.
- **(A3) Implicit Valid Transformation** An important implication of (A1) and (A2) is that once a specific transformation of input features into a composite score is established - typically based on one or more assumed outcomes - that same transformation is presumed to yield a representation that remains valid for predicting a broader range of distal outcomes. While rarely made explicit, this assumption is evident in the typical application of composite variables, where the same score is expected to carry predictive utility across different contexts or endpoints.

Many implementations of the theoretical model conform to a common statistical structure (**Figure 1**).^**4**^ This architecture consists of an input layer representing some observed features, a hidden layer that transforms these features, a second hidden layer containing a single node representing the composite score, and an output layer corresponding to one or more training targets assumed to be informative on the latent construct^**5**^. So-called “first-” and “second-generation” models can be viewed as specific variations of this architecture,

**Figure 1.**
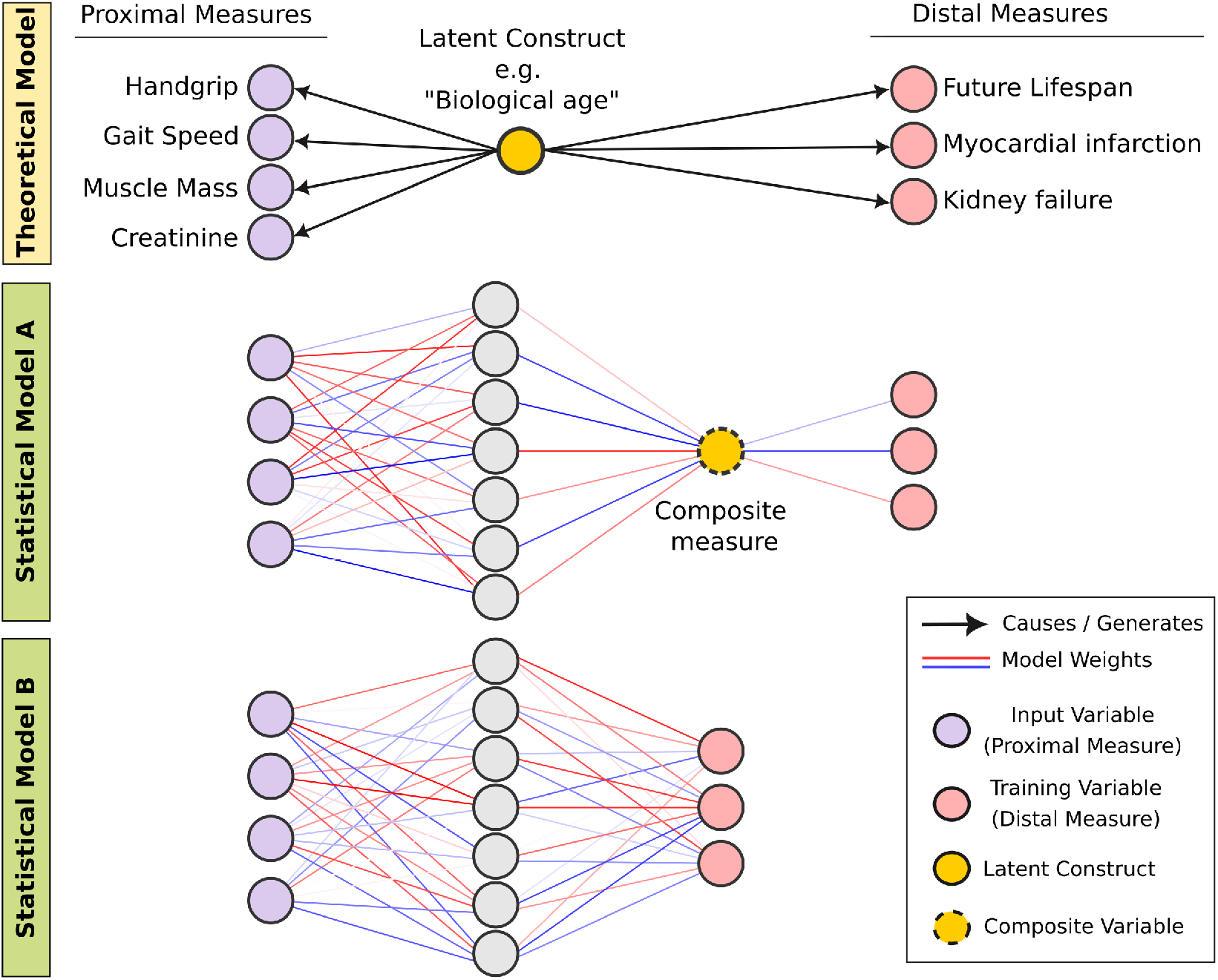
Conceptual and statistical models underlying composite “ageing” measures. (Top - Theoretical generative model) This model posits a latent construct - commonly referred to as “functional state” or “biological age” - which, though unobservable, is assumed to give rise to a set of measurable outcomes. The implication of this model is that more readily accessible measures can be used to estimate the latent construct, which in turn allows inference about distal outcomes - often the primary targets of interest. **(Middle - Statistical Model A)** Many implementations of the theoretical model conform to a common statistical structure. This architecture consists of an input layer representing some observed features, a hidden layer that transforms these features, a second hidden layer containing a single node representing the composite score, and an output layer corresponding to one or more training targets assumed to be informative on the latent construct. Line thickness and color reflect model weights; red and blue lines indicate positive and negative coefficients, respectively. **(Bottom - Statistical Model B)** A structural variant in which the intermediate composite layer is omitted. This represents the more appropriate reference model for evaluating the utility of composite ageing measures, as it tests whether the introduction of a latent construct - rather than simply relying on the same input features - adds value beyond what can already be achieved without such a representation.

with an identity function mapping the composite node to a single outcome node: chronological age in the “first generation”, and mortality risk in the “second”. While historically prevalent, this is only one of many possible instantiations of the generative model. Composite variables can also be derived using multiple outcomes [16]. In these cases, the model is not constrained to reproduce any single outcome, which weakens the common critique that a “perfect” model simply recapitulates its training target. Another common variation replaces the raw input features with their principal components, yielding an alternative, often decorrelated, representation of the original data [17]. Similarly, system-level or organ-specific “clocks” can be viewed as structural extensions of the base architecture described in Statistical Model A. The composite layer, instead of comprising a single node, consists of multiple nodes - each representing a distinct latent construct associated with a specific organ or physiological system. In these models, connections between input features and composite nodes may be selectively pruned or masked, such that only a subset of features contributes to each component score [8, 18].

Despite these differences, all these instantiations still share, to varying degrees, the same core structural assumptions outlined above. Assumptions A2 and A3 are often left implicit. In response to criticism, they are sometimes dismissed as non-issues, with claims that they can be “mitigated” - for example, by including raw features alongside the composite or by training separate composites for different outcomes. In contrast, A1 - feature convergence - is not optional. It applies to all statistical implementations of the latent construct, regardless of feature domain or modeling approach. It is a mathematically unavoidable property of collapsing multidimensional inputs into a lower-dimensional composite. This property is crucial to highlight because it directly contradicts one of the central motivations for developing composite “ageing” variables: the substantial variability in functional profiles observed among individuals of the same chronological age. Yet, by construction, these models inherit a similar limitation. This issue arises even in the simplest additive implementations with just a few input features where a large number of combinations can yield the same composite value (see **Figure 2** for a hypothetical illustration). This many-to-one mapping undermines the composite’s suitability for use as an exposure or outcome variable in many applications. These challenges have not gone unnoticed. Yet rather than prompting a reconsideration of whether such composites are suitable for certain objectives, the field has largely responded by doubling down. One strategy has been to build ever more “granular” composites - at the level of systems [19], organs [8, 18], tissue [20], cells [21], or even molecular pathways [22] - under the assumption that narrower scope reduces ambiguity. Another has been to fall back on the input features themselves in an attempt to “explain” the scores, typically through post-hoc attribution techniques marketed as explainable AI (XAI) [15, 23]. A third, increasingly common approach is to simply change the input space - by incorporating more features, higher-throughput assays, “emerging” technologies [24, 25], or by applying transformations or other embeddings [17]. A fourth approach has been to adopt trending “deep learning” models, under the assumption that increased model complexity and additional parameters can overcome the inherent limitations of these composite “age” variables for their intended applications as exposure or outcome measures [26, 27, 15].

**Figure 2.**
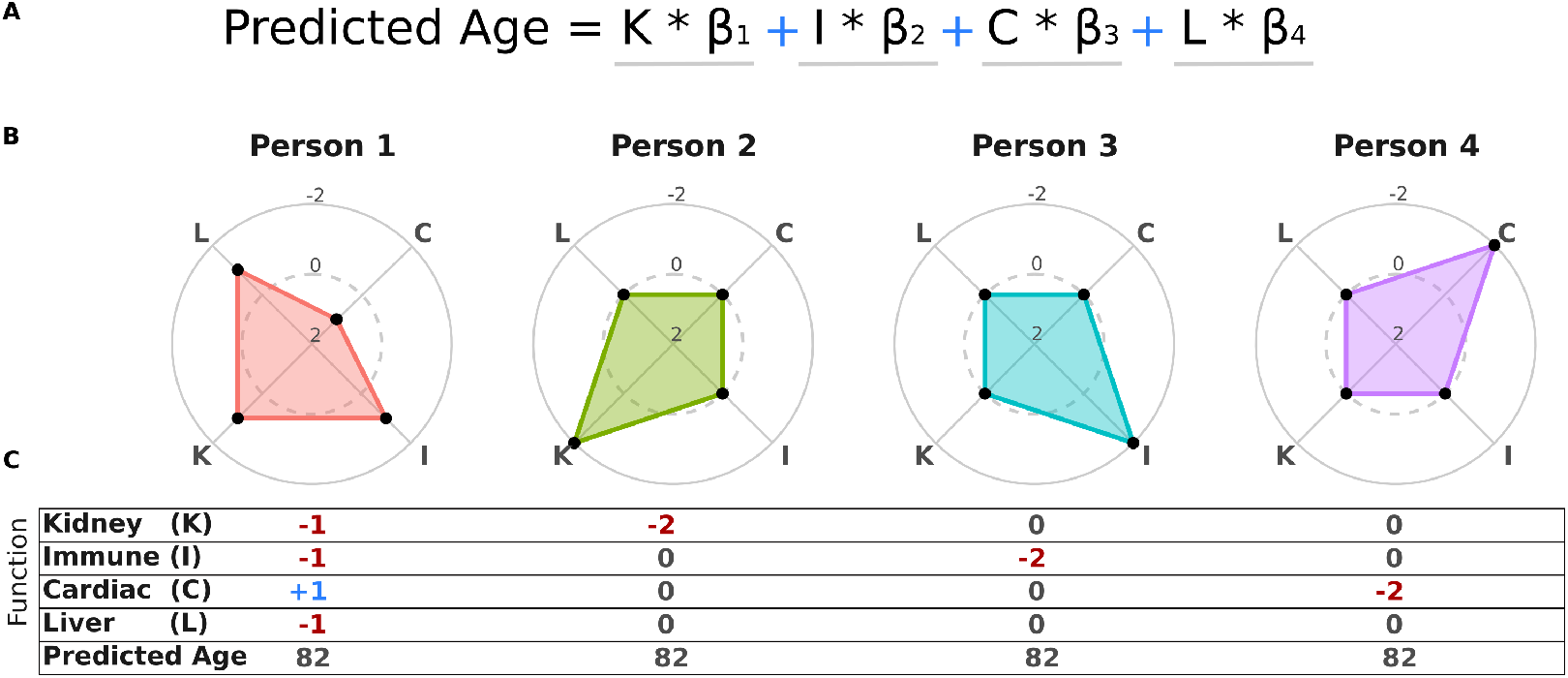
Hypothetical illustration of feature convergence (A1) in “ageing” Composites. **(A1)** Equation demonstrating the additive nature of predicted age as a weighted sum of functional contributions: Kidney (K), Immune (I), Cardiac (C), and Liver (L) functions, with respective weights (*β*_1_, *β*_2_, *β*_3_, *β*_4_). **(B)** Radar plots for four hypothetical individuals (Person 1–4) showing the z-scores of the four functional contributions. Each axis represents a function (K, I, C, L). The gray dashed circles mark z-scores of 0 (i.e., the average value for that function across all ages) and -2 (representing values two standard deviations below the mean). **(C)** Tabular representation of the z-scores for each functional contribution (K, I, C, L) for each individual, along with their predicted age. This example assumes equal weights (*β*_1_ = *β*_2_ = *β*_3_ = *β*_4_).

These models have been promoted for a wide range of clinical and research applications (**Table 1**). However, their added value is often assessed by comparing their performance to that of chronological age - for instance, by showing that the composite measure better predicts a given outcome. While this may suggest an improvement over a crude baseline, it sidesteps the more critical comparison: whether introducing an intermediate latent construct - such as “biological age” - adds value relative to using the same features within a model that does not rely on the assumption of such a representation (i.e., **Figure 1**, **“Statistical Model B”**, where the composite hidden layer is omitted). Importantly, the concern is not limited to predictive performance. The core issue is whether such a latent representation is helpful or even appropriate for the larger set of objectives these models are often intended to support, including intervention characterization, longitudinal monitoring, risk stratification, and mechanistic investigation (**Table S1**).

This article examines the structural implications of composite variables developed in the field of “ageing”, with particular emphasis on A1. First, using real-world data from the Baltimore Longitudinal Study of Aging (BLSA), we illustrate how the feature convergence assumption manifests in practice. Second, drawing on both cross-sectional and longitudinal data, we demonstrate how this property constrains monitoring and stratification. Third, we demonstrate that the same structural issues persist in molecular implementations - specifically in proteomic first- and second-generation “clocks”. Fourth, we examine their impact on objectives including intervention evaluation, outcome prediction, and investigation. Fifth, we discuss how these limitations are not specific to regression models. Finally, we consider additional properties that may help explain the widespread appeal and persistence of these models - despite clear conflicts with their stated aims.

## 2. Methods

### Composite Construction from BLSA Data

Multiple linear regression is among the most commonly used methods for constructing composite variables in “ageing” research [14, 15, 28]. These models are not only historically prevalent but also particularly well-suited for illustrating the convergence assumption - i.e., the Many-to-One Mapping property - due to their relatively simple additive structure. While the property can be demonstrated mathematically or using hypothetical data (Figure 1), its practical implications are most clearly revealed through real-world applications. To that end, we constructed sex-specific linear models to predict chronological age using data from the Baltimore Longitudinal Study of Aging (BLSA). The dataset was accessed through the Alzheimer’s Disease Data Initiative (ADDI) Workbench. A set of 27 features (**Table 2**) were selected to reflect cognitive, molecular, and vitality measurements. These included 12 cognitive variables (trats, trbts, bvrtot, crdrot, cvltca, cvlfrl, digfor, digbac, dsstot, boscor, flucat, flulet), 5 molecular variables (ldl, hdl, trigs, hgb, hba1c), and 10 vitality variables (handgrip, sbp, dbp, pe67hrt, r6mtime, gaitspeed, ftstime, slstime, cs5pace, nwalktime). These features were chosen to avoid redundancy and to exclude self-reported items or trivial scale transformations.

**Table 1.** Claimed applications of “ageing” composite metrics. Related to (**Table S1**).

| Category | Objectives |
| --- | --- |
| **Intervention** | - Trade-off Assessment<br>- Screening<br>- Development<br>- Monitoring<br>- Stratification / Personalization |
| **Investigation** | - Causal Mechanisms<br>- Individual Functional Variability<br>- Cross-species Variability<br>- Standardization |
| **Prediction** | - Risk Stratification<br>- Early Disease Screening<br>- Survival Prediction<br>- Clinical Trial Surrogate Endpoint<br>- etc. |

**Table 2.**
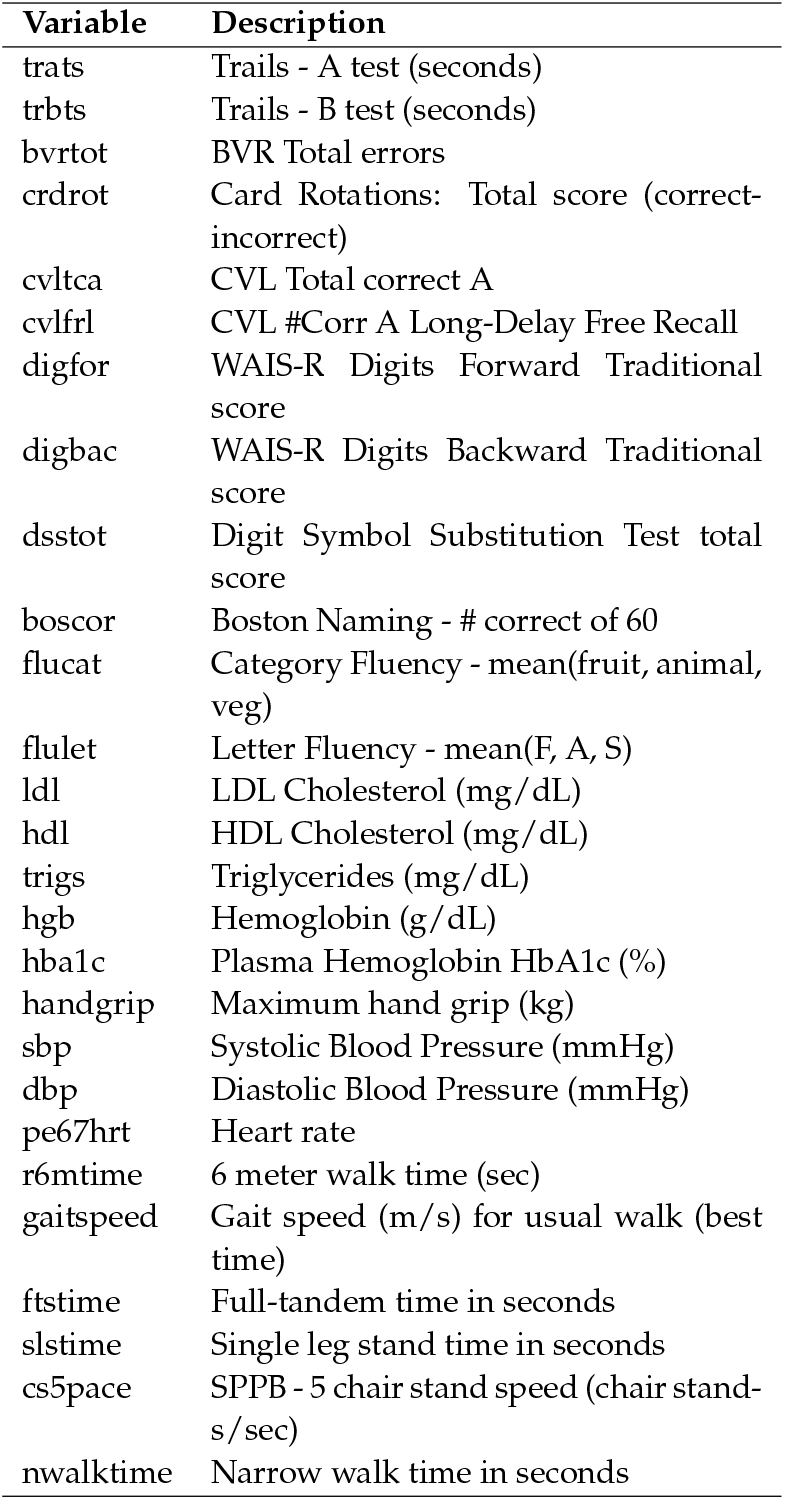
Descriptions of Selected BLSA Variables.

The dataset was subsetted by removing all rows with missing values for any of the selected features, as well as sex and age, resulting in a complete case analysis. The raw data included 3,495 patient visits corresponding to 753 unique participants. After restricting the dataset to complete cases over the selected features, 979 visits remained, representing 350 unique individuals (182 males and 168 females). For each sex, Z-scores were computed for each feature, using the mean and standard deviation across all individuals and ages. Two separate linear models—one for males and one for females—were trained using lm() in R with chronological age as the dependent variable and all 27 features as predictors.

To visualize the practical implications of the Many-to-One Mapping property, individuals whose predicted age (i.e. model output) was exactly 82 (after rounding) were extracted. Age 82 was chosen because it was the most frequently observed chronological age in the cohort (**Figure S2**). The z-scored input features of these individuals were plotted as a heatmap. Z-scores were computed across the entire cohort - across all individuals and all ages within each sex. The estimated model coefficients were visualized, stratified by sex, highlighting both sign and magnitude. To explore the stability in the composite for longitudinal monitoring, we identified individuals with at least three separate visits at which the predicted biological age was the same (after rounding), and plotted their z-scored input features across visits.

Another commonly used composite score in “ageing” research is “age deviation”, defined as the difference between an individual’s predicted criterion and the expected value given their chronological age. In the context of “first-generation” models - such as those trained to predict age - this corresponds to the difference between chronological age and predicted age. In “second-generation” models, which predict outcomes like mortality risk, age deviation reflects the discrepancy between predicted and expected risk at a given age. To illustrate how the Many-to-One Mapping property similarly affects this metric, we computed age deviation for all individuals and identified those whose deviation fell within a narrow window (4.9 to 5.0 years). The z-scored feature profiles of these individuals were visualized as a heatmap. Z-scores were computed across the entire cohort - across all individuals and all ages within each sex.^**6**^

### Molecular Composite Models from UK Biobank

To illustrate how these structural properties also apply to molecular “clocks” - particularly those built using high-dimensional “omics” data—we leveraged publicly available coefficients from a recently published set of proteomic clocks trained on UK Biobank plasma samples [18]. These models were constructed using elastic net regression and trained either on chronological age (“first-generation clocks”) or on mortality (“second-generation clocks”) as target outcomes. The original article used five-fold cross-validation and reported coefficient estimates separately for each fold. For the present analysis, coefficients from the first fold were selected to explore coefficient distributions. This fold was selected arbitrarily, as the observed patterns are not expected to vary meaningfully across folds.. All models were trained on a cohort of 44,952 individuals with proteomic profiles covering 2,916 plasma proteins, measured using the Olink Explore 3072 platform. Model weights used in this analysis were obtained from Supplementary Tables S1A (first-generation) and S1C (second-generation) of the original publication. For full details on model construction, and feature preprocessing, readers are referred to the original article. To examine the distribution and sign of model coefficients, we extracted the non-zero weights from the first cross-validation fold of both the first- and second-generation “conventional models” (i.e., models trained without organ-specific constraints). These weights were plotted as histograms to illustrate the extent of sparsity, symmetry, and magnitude of the estimated coefficients (Figure 4A and 4C). We then examined organ-specific implementations of both clock generations, using models trained on subsets of plasma proteins pre-defined as organ-enriched based on GTEx expression criteria (i.e., proteins with 4-fold higher expression in one organ relative to all others, as described in [8, 18]). For each organ-specific model, we computed the frequency of positive and negative coefficients. These were visualized as horizontal bar plots, stratified by organ (Figure 4B and 4D).

### Figures and Visualizations

All analyses and figures were generated in R version 4.4.1. The packages ggplot2 (v3.4.4), cowplot (v1.1.3), pheatmap (v1.0.12), reshape2 (v1.4.4), and tidyverse (v2.0.0) were used for visualization and data handling. Color palettes were specified using viridis (v0.6.5) and custom manual scales. Radar charts were implemented using a modified polar coordinate system. Neural network diagrams were generated using https://alexlenail.me/NN-SVG [29]. All figure panels were modified using Inkscape.

### Code and Data Availability

The code used to generate the simulated data and produce the figures, is available in the GitHub repository associated with this project (link). The BLSA Open Data utilized in this study is available through the Alzheimer’s Disease Data Initiative (ADDI) platform.

## 3. Results

### 3.1 BLSA Composite Models

To illustrate empirically the many-to-one property of these composites, we analyzed the Baltimore Longitudinal Study of Aging (BLSA) dataset. Linear regression models were constructed separately for males and females using a set of 27 features including cognitive, vitality and molecular features (**Figure 3A**). A detailed view of the functional profiles for individuals with predicted age of 82 is shown in (**Figure 3B**). As expected given the additive and compensatory nature of the model, various different combinations of feature values can result in the same predicted age label. For instance, some individuals may have high scores in some cognitive features offset by lower scores in some molecular measures, while others may show the opposite pattern. Furthermore, interpreting these profiles requires careful consideration of the coefficient estimates for each feature (**Figure 3C**). Some features have positive coefficients (e.g., HDL, where an increase in HDL leads to a higher predicted age), while others have negative coefficients (e.g., handgrip strength, where a decrease in handgrip strength results in a higher predicted age). Importantly, this interpretation extends beyond the sign or direction of the coefficients to their relative magnitudes. For instance, handgrip strength has approximately twice the impact on composite score value, with a coefficient of -2, compared to HDL, which has a coefficient of less than 1.

**Figure 3.**
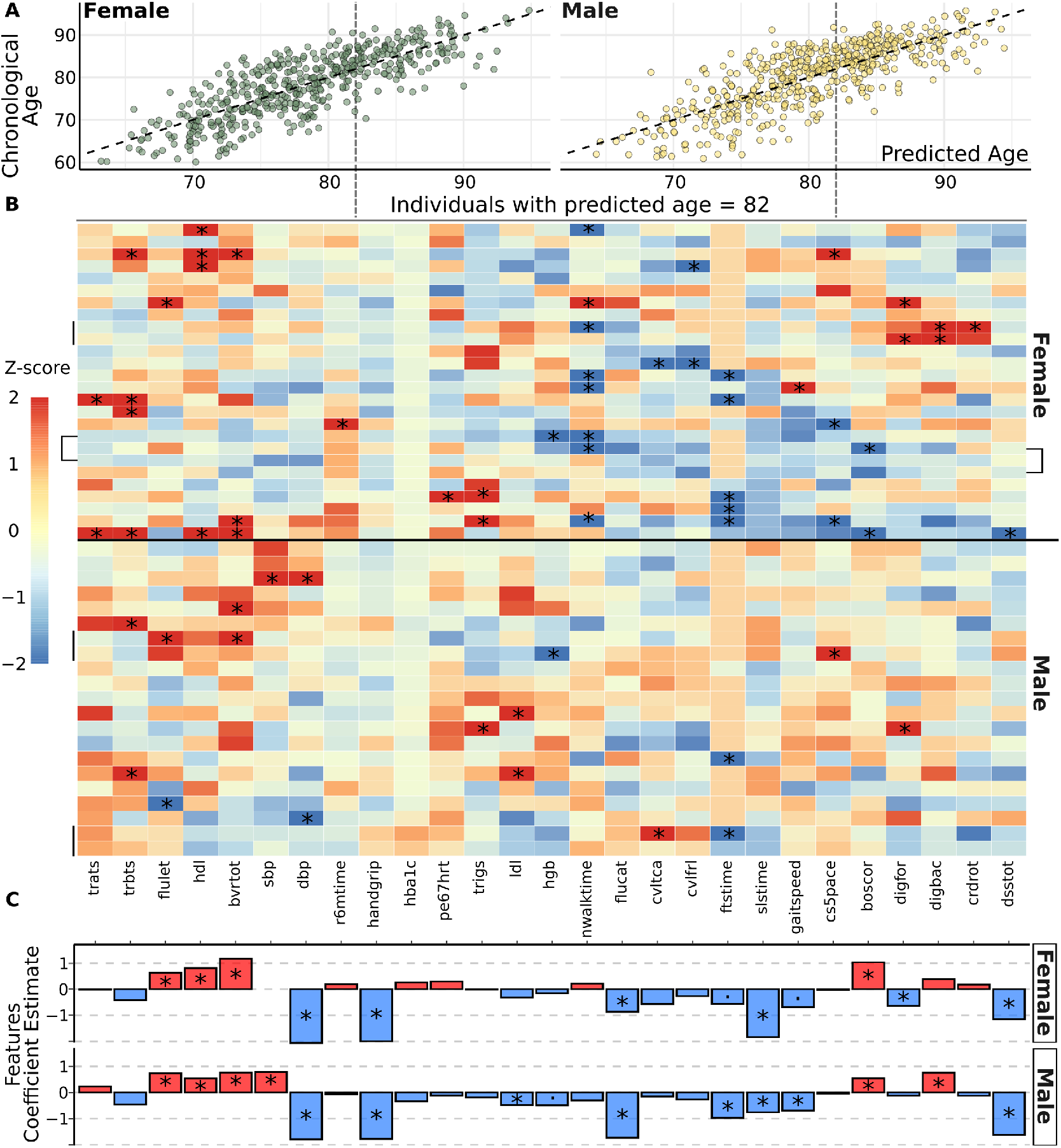
Cross-sectionally, distinct functional profiles can yield identical predicted age. **(A)** Scatterplots of predicted age (x-axis) versus chronological age (y-axis) for females (left panel) and males (right panel). Dashed black lines indicate the line of equality (predicted age = chronological age), and vertical dashed lines mark a predicted age of 82 years. **(B)** Heatmap showing z-scores (calculated per feature across samples of all ages stratified by sex) of features used to predict chronological age for individuals with a predicted age of 82. Columns represent features, and rows correspond to individuals. Red and blue cells indicate positive and negative z-scores, respectively, while black asterisks (*) highlight z-scores thresholded at -2 and +2. Vertical black lines indicate multiple visits from the same individual. **(C)** Standardized coefficient estimates for associations between input features and chronological age, stratified by sex (top: females; bottom: males). Blue bars indicate negative associations, while red bars indicate positive associations. Asterisks (*) denote statistical significance (p-value < 0.05) of the partial F-test (Supplemental Table S2-3).

**Figure 4.**
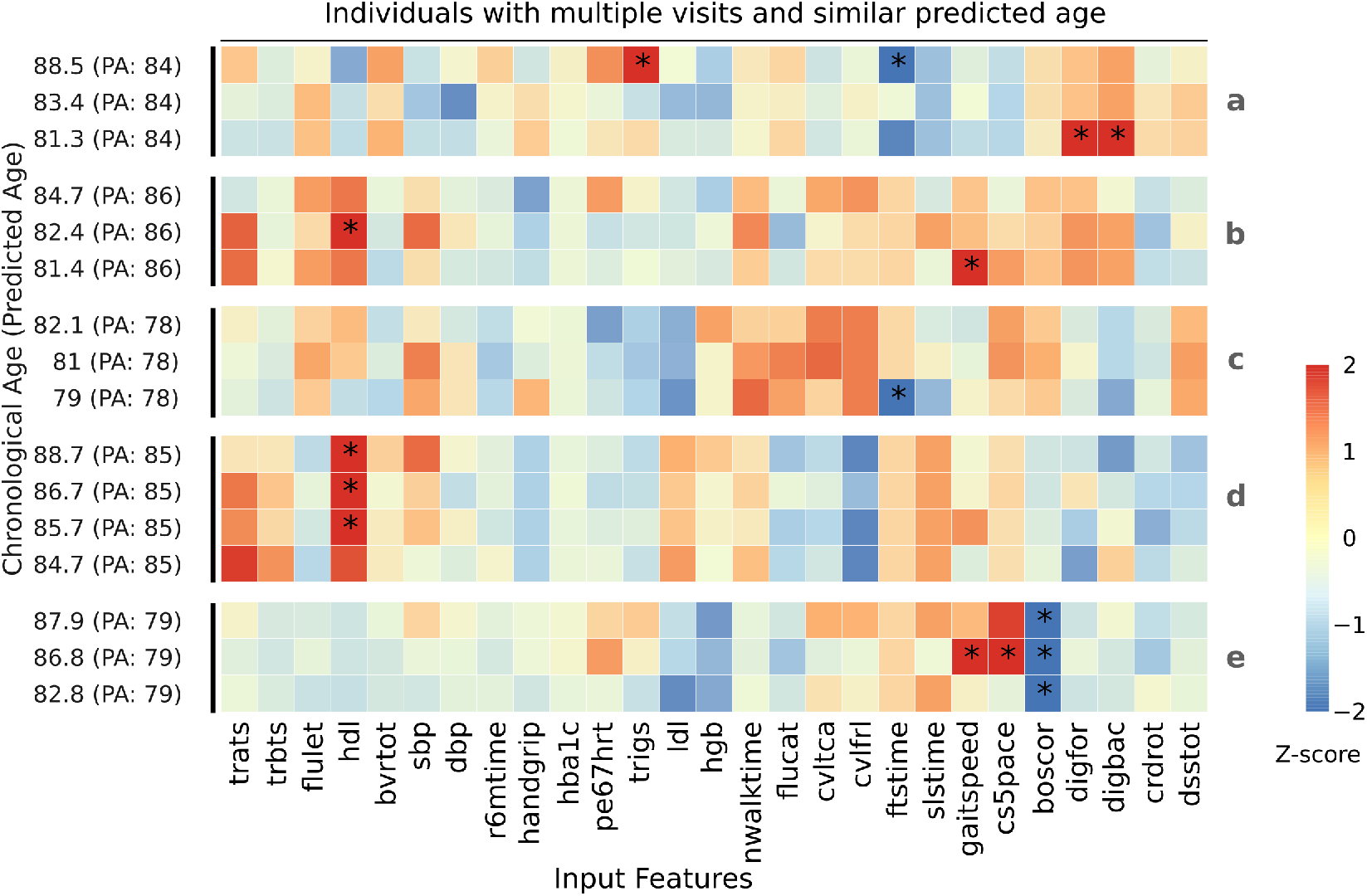
Longitudinally, predicted age can remain constant despite changes in functional profiles. Heatmap showing z-scores (calculated per feature across samples of all ages stratified by sex) for five individuals (a to e) with same predicted age across multiple visits. Rows correspond to individual visits, labeled with chronological age (CA) and predicted age (PA) in parentheses. Columns represent input features. Red and blue cells indicate positive and negative z-scores, respectively, while black asterisks (*) highlight z-scores thresholded at -2 and +2.

This challenge of reverse inference - starting with the composite score and attempting to deduce the underlying functional profile - also extends longitudinally for the same individuals. Examining individuals with multiple visits but a consistent predicted age label highlights this point (**Figure 4**). For example, Individual (**a**) is consistently predicted to have an age score of 84 across three visits spanning seven years. However, their underlying functional features did not remain as stable. In contrast, Individual (**d**) shows relatively stable feature profiles across visits, with minimal changes in cognitive, vitality, or molecular measures. These examples illustrate the implications of the additive property of these models in the context of patient follow-up: a consistent predicted age across multiple visits does not necessarily indicate stability in the underlying functional features. As a summary measure, “composite age” can obscure significant changes within the multidimensional space of contributing features, limiting its utility for tracking individual health trajectories without fallback on its components.

Another widely employed composite metric is “age deviation” - defined as the difference between an individual’s predicted criterion and the expected value of that criterion, sometimes adjusted via regression on additional covariates. This measure is frequently interpreted as capturing “how fast aging processes are occurring at a given point in time” [10], and is commonly used in “first-generation” (age-predicting) and “second-generation” (mortality-predicting) models (REF). Besides the broad historical criticisms of this metric, it suffers the same challenges for applications as the “age” composite scores. Even if one assumes this quantity to be “meaningful”, the same deviation can be created by various combinations of features depending on the feature covariance and model structure. To illustrate this, we identified individuals from the BLSA dataset with “age deviations” tightly clustered between +4.9 and +5.0 years (**Figure 5A**). Despite this shared deviation value, the individuals exhibited highly heterogeneous profiles (**Figure 5B**).

**Figure 5.**
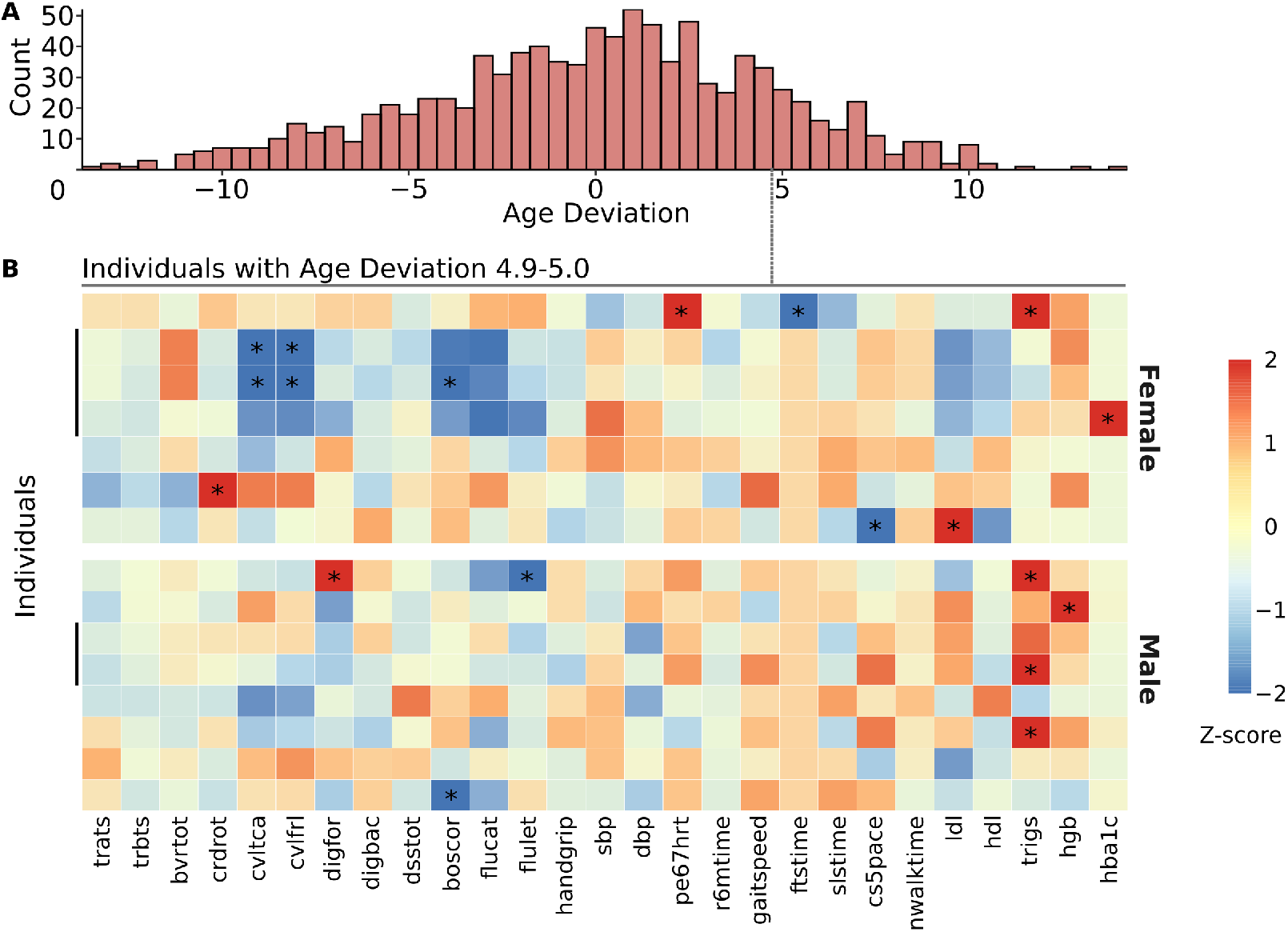
Heterogeneity among individuals sharing the same +5y age-deviation label. **(A)** Distribution of age deviation in the BLSA cohort. Age deviation was calculated as the difference between each visit’s predicted age (output of sex-specific linear models trained on 27 features) and the corresponding chronological age. The histogram uses a bin width of 0.5 years. The grey dashed line at +5 years marks the age deviation window (4.9–5.0 years) corresponding to the individuals shown in panel B. **(B)** Heterogeneity among individuals sharing the same +5y age-deviation label. Heatmap showing z-scores (calculated per feature across samples of all ages stratified by sex) for 15 individuals (7 female, 8 male) with same +5y age-deviation. Rows correspond to individual visits. Columns represent input features. Red and blue cells indicate positive and negative z-scores, respectively, while black asterisks (*) highlight z-scores thresholded at -2 and +2. Vertical black lines indicate multiple visits from the same individual.

### 3.2 Molecular Composite Models

These properties and challenges are more pronounced in molecular composite measures of “ageing”, particularly those built from high-dimensional “omics” data. These models typically rely on hundreds of features, increasing the number of distinct input profiles that can produce the same composite score. While this number also depends on the covariance structure of the features, the coefficient distribution provides a useful indication of how diverse those profiles can be. For the “first-generation model”, 2,315 non-zero coefficients were retained, ranging from -1.16 to 1.65 (1st quartile = -0.047, mean = 0.0003, 3rd quartile = 0.039) (**Figure 6A**). For the “second-generation model”, 148 coefficients were retained, ranging from -0.20 to 0.35 (1st quartile = -0.015, mean = 0.0052, 3rd quartile = 0.020) (**Figure 6C**). In both models, the distributions are centered near zero and span both positive and negative domains, reflecting heterogeneous directions of association between features and the composite score. This spread of weights - in both magnitude and sign - underscores how distinct combinations of protein levels can yield the same composite score.

The ambiguity inherent in these composite scores has not gone unnoticed. One response has been the development of more “granular scores” - e.g. organ-level models. In the “first-generation clocks”, the number of retained features ranged from 123 for the immune model to just 2 for the thyroid (**Figure 6B**). In the “second-generation clocks”, the range spanned from 85 (immune) to 2 (thyroid) (**Figure 6D**). Most organ models showed a mix of positive and negative coefficients, though polarity became more skewed in models with very few features. While these lower-dimensional models might appear to offer more “interpretable profiles” - and in principle reduce the number of input configurations compatible with a given set of composite scores - they still inherit the same property. A given “heart age” score, whether from a first- or second-generation model, may correspond to a wide variety of underlying proteomic states. In turn, these divergent profiles could imply very different intervention targets, despite producing the same composite score.

**Figure 6.**
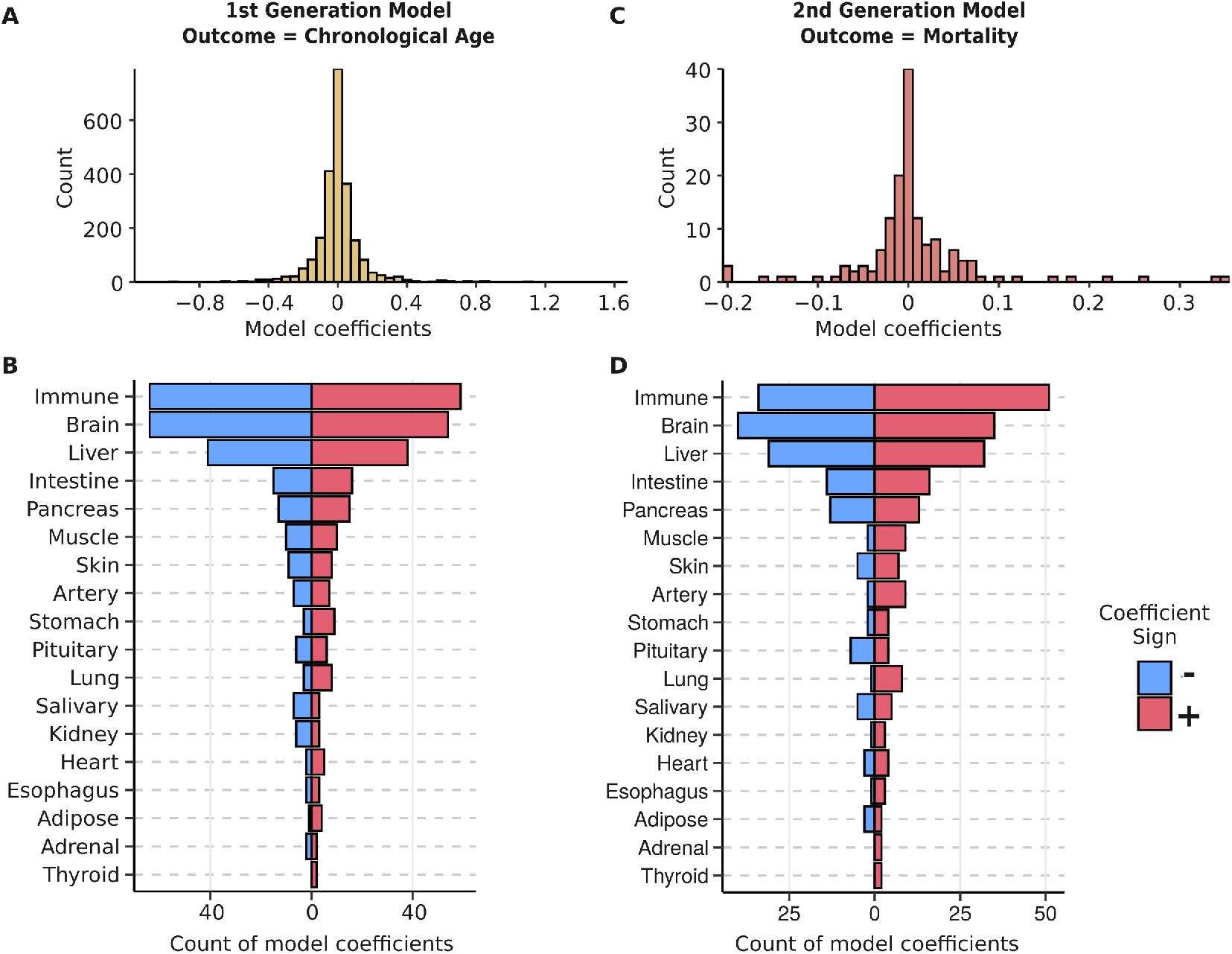
Feature convergence (A1) in molecular “ageing” composites. **(A)** Histogram showing the distribution of non-zero model coefficients from a “first-generation” elastic net model trained to predict chronological age using plasma proteomic data (n = 2,315 features). **(B)** Bar plot showing the number of retained features with positive (red) and negative (blue) coefficients for each “first-generation organ-specific” model. **(C)** Histogram showing the distribution of non-zero model coefficients from a “second-generation” elastic net model trained to predict mortality risk using the same proteomic dataset (n = 148 features). **(D)** Bar plot showing the number of retained features with positive (red) and negative (blue) coefficients for each “second-generation organ-specific” model.

## 4. Discussion

The literature surrounding composite “ageing” measures is often mired in rhetorical claims on how these metrics reflect “the health status of an individual” [23] or “the very mechanisms of aging” [30]. Such claims offer little utility as the purported object of that reflection - a latent construct - is, by definition, unobservable. Consequently, debates over construct validity often devolve into circular reasoning and validity by assumption. Yet, validity and reliability of a model are not intrinsic properties; they emerge only in relation to a specific purpose. Chronological age could be said to “reflect ageing”, is predictive of mortality, yet no one would seriously propose it as a suitable metric for patient monitoring. Accordingly, the present discussion does not dwell on abstract considerations of construct validity. Instead, it will focus on the added utility of these composite scores for specific categories of objectives: intervention, prediction, and investigation.^**7**^ Before examining each objective, it is important to clarify a few foundational considerations. First, the key comparison is not between using a composite measure and collecting no data, nor between the composite and chronological age. Rather, it lies between the composite measure and its features - specifically, between a model that assumes a latent construct (**Statistical Model A**) and one that uses the same features without invoking such a latent representation (**Statistical Model B**). The central question - often overlooked - is whether the composite offers added value beyond its inputs. Second, throughout the following sections, reference will be made to three key properties of composite measures:

- **(P1) Simplicity**: These constructs serve as summary statistics. By reducing a multidimensional feature space to a single value, they simplify reporting, communication, and comparison between individuals or groups.
- **(P2) Pooling**: Useful when decisions or outcomes depend on the combined contribution of multiple variables rather than any one in isolation. Including correlated features makes the composite less sensitive to fluctuations in individual variables. This redundancy improves stability, reduces the impact of noise, and increases detection power. When multiple correlated features shift together, the composite can offer stronger evidence of systematic change, especially when effect sizes are small. Conversely, composites can also be structured so that a change in a single component - while others remain constant - is enough to shift the overall value.
- **(P3) Many-to-One Mapping**: As outlined in A1, different combinations of feature values can yield the same composite score. This undermines direct backward inference. Knowing a composite value often reveals little about the specific configuration of the underlying variables.^**8**^

Third, as the magnitude of these properties is a function of the covariance of the composite’s input features, evaluating their utility requires considering two theoretical extremes. Real-world models typically fall somewhere between these extremes, combining characteristics - and limitations - of both.

- **(E1) Perfect Correlation Between Features**: If the features are highly correlated, the model exhibits strong redundancy, making it robust to fluctuations in any single feature and less sensitive to measurement error. Additionally, these models remain highly interpretable, as any increase or decrease directly reflects a proportional shift in all features. This is a result of the rigid covariance structure reducing the (P3) property, as different feature combinations are far less likely to produce the same value compared to a scenario where features are weakly correlated. Thus, in this scenario, (P1) does not come at a high cost of lost information.
- **(E2) No Correlation Between Features**: If the features used are weakly correlated, the model exhibits strong (P3), meaning that vastly different feature combinations can produce the same composite value. As such, (P1) comes at a high cost of lost information, making it difficult to interpret. This issue becomes evident when attempting to interpret an increase in the composite. Such a change could result from an increase in some features while others remain constant, or from a large increase in certain features alongside a smaller decrease in others. Distinguishing between these scenarios is critical for certain objectives, as will be discuss. Moreover, no change in the composite cannot be interpreted as stability at the feature level, since opposing shifts among uncorrelated features may cancel each other out.

Finally, many of the limitations discussed apply across multiple objectives. To avoid redundancy, such issues will be addressed only once where most relevant.

### 4.1 Intervention

One of the key underlying goals behind employing composite “ageing” variables is better intervention characterization. This is most evident in the long-standing criticism of using lifespan as the sole measure of intervention success, where improvements of this measure may come at the expense of function [31]. As any intervention comes at a cost, the critical task is to identify this cost and evaluate whether the benefits justify it. The problem is that these trade-offs are not easily predictable due to the system’s complexity [32]. Unintended consequences can arise in numerous, often unforeseen, ways. To identify these trade-offs, one must observe the system for a “sufficient” duration and with “enough detail”; otherwise, crucial effects might be missed. While more information is generally useful, collecting it always comes at a price. Every additional variable measured comes at a cost to the limited resources available - from funding to time. Especially in long-term studies, the more data we attempt to gather, the greater the demands on resources, time, and logistical feasibility. As such, one has to prioritize a set of measurements to focus on otherwise it is unfeasible. “Ageing” composites are positioned as tools that address this need. Proponents claim they improve intervention characterization, allowing for the detection of potential trade-offs, facilitate screenings, and guide the development process itself (**Table S1**). Additionally, they are presented as tools for intervention personalization and stratification, allowing for the identification of individuals most likely to respond to specific interventions.

Lets consider each of these objectives in that order. First, regarding intervention characterization and the detection of trade-offs. In the E1 scenario (highly correlated features), unforeseen trade-offs are inherently absent, as all features shift together. In the E2 scenario (weakly correlated features), P4: Many-to-One Mapping causes the composite to obscure rather than reveal trade-offs. An increase in the composite could result from an increase in some features while others remain constant, or from a large increase in certain features alongside a smaller decrease in others - i.e., a trade-off. Distinguishing these cases requires returning to the feature space. Thus, the composite offers no added value over raw features for detecting trade-offs, neither improving their identification nor simplifying interpretation.

Second, the primary goal of screening is to quickly and cost-effectively prioritize potentially promising interventions for more comprehensive evaluation. Given the large number of interventions typically tested, speed and minimal resource investment are key considerations. Composite measures do not reduce measurement costs, since they require assessing all component features. Instead, their primary advantage is simplifying analysis, as comparing a single value across treatment and control groups is undeniably easier than working with multiple features. Considering the two extreme cases, in the E1 scenario, any effective intervention would proportionally shift all features. Here, using a composite contradicts the goal of simplicity - directly comparing just one or two representative features would yield the same conclusion. In the E2 scenario, the composite appears more useful but introduces an additional assumption: only interventions that shift certain features more than others will be detected. Since a stable composite does not necessarily indicate intervention failure, meaningful changes at the feature level could be overlooked. To avoid this, one would still need to analyze the feature space. Overall, while a composite might provide some added value as a quick comparison tool for screening, the magnitude of this benefit may be modest compared to simpler, rapidly computed statistics derived directly from the underlying features.

Third, for monitoring, the E1 scenario presents a fundamental contradiction to the objective, as effective monitoring inherently requires measuring features that are not highly correlated to capture a comprehensive and nuanced picture of an individual’s health state. In the E2 scenario, a different contradiction emerges: while it aligns with the goal of capturing diverse and independent aspects of health, the use of a composite in this context undermines both cross-sectional and longitudinal monitoring. Cross-sectionally, individuals with the same composite can have different underlying feature profiles, as demonstrated by the BLSA dataset (**Figure 3**) and the hypothetical example (**Figure 2**). Longitudinally, a stable composite across multiple visits does not imply feature stability, as individuals with unchanged labels over time can still experience substantial shifts in their individual features (**Figure 4**). Without directly analyzing the feature space, these changes remain undetected, defeating the core purpose of monitoring health trajectories.

Finally, these same contradictions extend to intervention stratification and personalization. The stated goal of both objectives is to enable individual-specific profiling, yet the composite’s limitations undermine this aim. The E1 scenario inherently contradicts the goal of personalization and stratification, which rely on measuring distinct aspects of health to tailor interventions. Conversely, in the E2 scenario, the composite becomes even less informative. Two individuals with the same score can have vastly different underlying health profiles, rendering the composite measure inadequate for personalized intervention design. For example, targeting kidney function might benefit an individual whose an elevated score is driven by renal decline, but the same intervention could be irrelevant - or even harmful - for another individual with the same score driven primarily by cognitive impairment. In both cases, reliance on the composite not only fails to advance stratification and personalization but risks misleading conclusions by masking critical individual-level variability. This underscores the necessity of returning to the feature space to capture the nuanced information required for effective intervention tailoring, reducing the composite “ageing” scores to nothing more than a superficial and uninformative summary statistic.

### 4.2 Prediction

One might argue that, even if composite measures offer little added value for intervention characterization, they could still hold value in predictive contexts. For such objectives, the priority is not to understand how the prediction is generated but simply to forecast future outcomes with greater validity. Some proponents argue that these composites improve validity, reliability, and reduce overfitting - benefits expected to help in both research and clinical settings.

First, on improved prediction validity. The core issue lies in the comparison underpinning such claims. The added predictive value of these composites is often framed against chronological age or a minimal set of standard clinical measures. But the relevant comparison is between the composite measure and its features - specifically, between a model that assumes a latent construct (**Statistical Model A**) and one that uses the same features without invoking such a latent representation (**Statistical Model B**). By aggregating multiple variables, the composite measure adds an extra layer of abstraction and can discard valuable information. If the original features are truly predictive, using them directly will always be at least as effective - if not superior - than compressing them into a single score. This is especially important when predicting multiple outcomes. Collapsing diverse features into a single value assumes a fixed relationship among them, regardless of the outcome. Reality is rarely that uniform [33]; feature relevance and interactions may shift depending on the outcome being modeled - be it mortality, cognitive decline, or cardiovascular events. This contradiction is clearest when “deep learning techniques” - designed to capture non-linear feature interactions and complex relationships - are used merely to generate rigid composite scores [15].

Second, the same applies to claims of improved reliability and noise reduction. Any noise reduction attributed to these composite measures stems from the aggregation of correlated features, not from the composite itself. If the goal is prediction, directly modeling the outcome from these correlated features inherently captures the same benefit. In addition, as was the case for validity when multiple outcomes are considered, collapsing features into a single score imposes a fixed noise structure that may not align with the specific noise characteristics of different predictions.

Third, the notion that composite-based models are “simpler” and therefore less prone to over-fitting is often overstated - especially when only one outcome is being predicted. When applied in such a context, Statistical Model A in Figure 1 (with a single distal measure) would contain virtually the same parameter count as Statistical Model B: the weights that generate the composite in Model A are simply redeployed in Model B to map directly from the hidden layer to the outcome. Complexity is not reduced; it is merely hidden behind the façade of a latent score. With multiple outcomes, the claim has slightly more substance: omitting the final set of connections from the hidden layer to each outcome in Model A saves roughly (hidden-layer nodes × number of outcomes) parameters. Yet this saving is modest relative to the size of the full network and comes at a price - forcing all outcomes to depend on the same latent representation, which can hamper flexibility and predictive accuracy. If over-fitting is the genuine worry, established strategies such as regularisation, cross-validation, feature selection, training-validation splits with a held-out test set, and dimension-reduction techniques can help deal with model complexity directly, without the strong structural assumptions imposed by a composite measure.

### 4.3 Investigation

Investigations aim to uncover the generative processes behind variability - that is, the causes [16]. A practical definition of a cause is its ability to predict the outcome of an intervention that alters it. Investigations search for potential causal variables, which are then tested through interventions. Relying on “ageing” composite measures for either objective can be misleading and counterproductive. First, these composite measures are non-actionable. Because they cannot be directly manipulated, any observed associations risk misinterpretation without specifying the underlying intervention [34, 35]. The outcome is determined by the specific intervention used to change the composite measure - not by the change in the composite measure itself. A case study later in the article will examine this issue in greater depth. Second, relying on these composites weakens hypothesis generation. Different underlying processes can produce the same composite value through fundamentally distinct mechanisms. This issue is amplified in cross-species studies, where dissimilar features are collapsed into superficially similar composite measures, leading to false equivalences and misleading conclusions^**9**^. Third, these composite measures often reinforce their own built-in assumptions rather than generating new “insights”. Recognizing these assumptions is crucial, as they heavily shape findings. This circularity is most evident in how such measures are developed and validated. For instance, if a composite is designed to distinguish calorie restriction as an “anti-ageing” intervention, it will naturally succeed in that context. But when applied to assess something unrelated - like radiation exposure or some hypothesized “anti-ageing” drug - ambiguous results trigger experimenter’s regress [32, 37]: is the intervention ineffective, or is the measure simply not adequate for this objective? Fourth, proponents argue that such composite measures improve standardization by providing a unified metric for comparing studies, populations, and methods. Because such composites require measuring all its components, it is thought to reduce variability in data collection and facilitate meta-analyses, reproducibility, and generalization. But if the goal is consistent measurement, directly specifying required variables achieves this without the added abstraction and complications of a composite score. In practice, these measures have done the opposite of standardizing the field: rather than converging, they have splintered into inconsistent definitions, variable sets, and methods. As a result, “ageing” composites often aren’t comparable across studies, introducing confusion rather than clarity. Useful standardization comes from aligning what is measured, not from collapsing diverse inputs into an abstract, inconsistent composite.

### 4.4 Case Study: Body Mass Index (BMI)

The Body Mass Index (BMI) is an instructive case study of the risks of using composite measures for certain objectives. Though far simpler than “modern” biological age (BA) composites, it shares many of the same properties and limitations. It is an ideal example for three reasons. First, its popularity stemmed from its use as a crude and accessible proxy for body fat - much like “ageing” composites are positioned as proxies for “biological age” or “ageing state”. Second, its broad use in research, clinical practice, and public health parallels intended uses of BA composites. Third, its simplicity and the intuitive nature of its features makes the properties - often obscured in BA composites - more obvious. For example, the Many-to-One Mapping property (P3) is particularly intuitive in the case of BMI: multiple combinations of height and weight can produce the same BMI value. Furthermore, even among individuals with the same height and weight, BMI inherits a fundamental limitation from its reliance on weight. An athlete with high muscle mass and a sedentary individual with muscle loss and excess fat could have the same BMI, yet their health risks would be drastically different. Furthermore, even if two individuals had the same weight and the same proportion of fat, the type and distribution of that fat might still matter.^**10**^

The impact of this property became evident when BMI was used to monitor interventions. A change in BMI can result from very different shifts in body composition and provides no indication of what changed - fat, muscle, or both. Two individuals may show identical reductions in BMI yet experience sharply different health outcomes depending on what was lost [38]. This ambiguity is problematic in research but is even more consequential in clinical settings. This is because the change in BMI itself is not the primary interest but the interpretation of that change in terms of future health risks. This problem became even more pronounced when BMI was treated as an exposure rather than just an endpoint. A central issue in causal inference is that BMI is not an actionable variable - it cannot be directly manipulated [38, 39, 40]. Instead, specific interventions have to be used, such as diet, exercise, or pharmacological treatments. For example, lowering BMI through calorie restriction alone may reduce both fat and muscle, potentially increasing frailty. In contrast, combining diet with resistance training could preserve or even build muscle mass. Similarly, some medical conditions reduce BMI but worsen health outcomes. Despite the same numerical BMI change, these different interventions can be associated with vastly different health outcomes. This makes BMI a textbook case of an ill-defined intervention. Without knowing what caused the change, it can be misinterpreted - as reflected in the different associations between BMI and mortality reported across studies [41]. Ultimately, the solution is simple: rather than asking “What is the effect of lowering BMI on an outcome e.g. mortality?” the question should be “What is the effect of a specific intervention - such as one hour of resistance training - on mortality?” This reframing shifts the emphasis to what actually drives outcomes - the intervention - rather than a low-resolution, non-actionable and hard-to-interpret proxy. This limits misinterpretation, avoids treating BMI as explanatory, and keeps the focus on actionable conclusions. Finally, when BMI is used for prediction tasks, it imposes critical assumptions. By collapsing height and weight into a single metric, it assumes a fixed relationship between them - regardless of the outcome being modeled. Yet this relationship likely varies by context and health outcome. For example, Sorjonen et al. [42] analyzed data from 49,000 Swedish men and found that “BMI is not an optimal way to model effects of weight and height on mortality. In this sample, a model with a linear effect of height and a quadratic effect of weight was superior”. In practical terms, BMI imposes an arbitrary constraint on the relationship between height and weight, limiting a model’s ability to capture context-specific associations. Using height and weight directly improves predictive power and model flexibility.

“Ageing” composite measures share these same limitations - only to an exacerbated degree. First, both serve as proxies for another variable of interest. For BMI, the target is adiposity; for BA composites, it is the concept of “biological age” or “ageing state”. A key difference, however, is that adiposity is relatively well-defined and measurable, whereas “biological age” remains a nebulous and poorly defined concept [32, 43, 44, 45, 46, 47]. Yet even with this simplicity, BMI has faced persistent challenges in interpretation and has often proven to be a poor choice as both an exposure and an outcome in intervention studies [38, 39, 40]. The conceptual ambiguity surrounding BA only amplifies these issues, making its limitations more severe.^**11**^ Second, BMI is derived from just two variables - height and weight - both of which are intuitive and, in the case of height, largely stable in adults over time. Yet, even with such a formulation, BMI has struggled with issues of misinterpretation and improper use as an exposure [41]. In contrast, BA models incorporate dozens to hundreds of variables, often relying on far more complex and opaque model architectures [18, 35, 27]. This dramatically worsens the many-to-one mapping problem, multiplying the possible combinations that produce the same composite score and further obscuring interpretation. Third, like BMI, BA composites are not actionable variables - they cannot be directly modified. Given the diversity of input features, there are far more ways these composites can change compared to BMI. Two individuals could show the same reduction in a BA composite driven by entirely different changes - just as identical BMI reductions can reflect very different shifts in body composition. This greater complexity exacerbates the interpretive uncertainty.^**12**^ Finally, for prediction, BA models face the same problem as BMI: they collapse diverse variables into a single score, assuming a fixed relationship between them regardless of the outcome. This risks discarding valuable information and reducing predictive utility, especially when individual components have nonlinear or context-specific effects.

### 4.5 Modeling and Empirical Discovery

In “ageing” research, it is common to construct composite metrics, suggest their potential utility for various objectives, and call for future validation or causal investigation to provide definitive answers [43, 48]. Although empirical validation is indispensable, it is not always required to recognize when a model is structurally unsuited to a given task. A major theme in this work is that many pitfalls of “ageing” composites arise not from surprising empirical results, but from their underlying structure. Once a model’s assumptions and architecture are made explicit, many so-called “findings” become predictable. Perhaps the most elementary example of this is the widely “discovered” limitations of so-called “first-generation” composites. This is illustrated in a recent perspective (Moqri et al., 2023), which states: “a recent study [49] suggested that when the performance of a biomarker of chronological age approaches near-perfect predictive accuracy, its association with mortality attenuates”. One did not need to wait for empirical discovery, new data, or improved methods to anticipate this outcome - it follows directly from the structure and logic of the model itself. In fact, this was clearly articulated over 45 years ago in “Functional Age: A Conceptual and Empirical Critique” [3]: “If the regression succeeded perfectly, the resultant statistical age should correlate 1.0 with chronological age, and hence would be a perfect and perfectly useless, alternative to chronological age”. This pattern has not subsided. The likely outcomes of many current trends in the development of “ageing” composites can already be anticipated from the outset.

One direction that has gained traction is the development of ever more “granular” composites - at the level of systems [19], organs [8, 18], tissue [20], cells [21], or even molecular pathways [22]. This trend is driven by the observation of heterogeneity among subscores constructed by partitioning a broader composite. Yet this “discovery” of subscore heterogeneity is not surprising. It is a direct and inevitable result of constructing the original composite by selecting minimally correlated features [18, 28] and projecting them into a single score. Though narrower, these “granular” composites still carry the same property. A heart-specific metric, for instance, might seem more targeted than a whole-body metric, it still fails in the same way: two individuals with identical “poor heart scores” may have vastly different underlying conditions requiring entirely different interventions. The metric remains non-actionable and non-interpretable. Its predictive power, as before, stems from the component features rather than any added value of the composite measure. Ironically, if this trend continues, it will eventually come full circle - fragmentation will lead to the recreation of the original feature set, just under different names. Eliminating such heterogeneity would require either using highly correlated features or explicitly identifying which forms of variability are irrelevant to a given objective^**13**^. But doing so would undermine the generality and broad applicability that these scores are often claimed to offer. This relates to another “discovery”: that models trained to predict specific outcomes - such as cause-specific mortality - often outperform those targeting broader outcomes like all-cause mortality [15]. Unless one assumes, as the latent construct model does, that a single composite meaningfully captures all relevant variation across diverse outcomes, there is little reason to expect a general model to outperform a task-specific one.^**14**^

Another popular trend is the creation of so-called “next-generation” composites, either by modifying the input features or by adopting more fashionable modeling techniques. In some cases, the same modeling approach is applied to a different set of variables - often drawn from the latest high-throughput or “cutting-edge” technologies to lend a sense of novelty. In others, the feature set remains unchanged, but the model is swapped out for whatever “machine learning” method is currently in vogue - or both are altered at once. Yet whether one replaces linear regression with “deep learning”^**15**^ or expands a neural network from 10 to 100 layers, both approaches ultimately just compress the same feature set into a single output. More computation does not equate to a better composite metric, and increasing model complexity only exacerbates - rather than resolves - the structural limitations discussed so far. These newly constructed models are often accompanied by yet another recurring “discovery”: the observation that composites built from different input features, or trained on different outcomes, are said to reflect “different aspects of ageing”. But this only appears noteworthy under the assumption that all such composites should converge on a single underlying process. Once that assumption is relaxed, there is nothing surprising about divergent outputs; the models are, by design, built on different data and optimized for different objectives. Moreover, because the latent construct these models are meant to approximate is unobservable, nearly any empirical outcome can be retroactively interpreted to fit the theory. If two models diverge, it is said they capture different “aspects” of the latent construct - after all, no one claimed it was unidimensional. If they converge, it is taken as confirmation that they reflect the same underlying entity. This flexibility underscores the degrees of freedom such latent constructs afford, specifically, in the selective adoption of assumptions.^**16**^

Another trend is the search for “causal” versions or interpretations of these composites - either by explicitly constructing them around assumed biological mechanisms, or by probing how those mechanisms might drive changes in the score [15]. Yet neither strategy overcomes the fundamental drawback that makes such composites poor choices as outcomes or exposures. As the composite is non-actionable, any change must be produced through a concrete intervention (e.g., diet, exercise, drug treatment), and the biological meaning of that change depends entirely on which intervention was used. This intervention-dependence leaves the composite ill-defined for causal questions, because the quantity “change in the score” has no uniform interpretation across different experimental or clinical contexts. As such making causal claims based solely on changes in the composite are ambiguous at best, and misleading at worst. Yet instead of reconsidering whether such composites are suitable exposures or outcome variables, the field has pivoted toward a new sub-trend marketed as “explainableAI” [23]. Under this banner, various metrics - feature importances, saliency maps, Shapley values - are employed in an effort to “explain the score” to “potentially guide clinical decision making” [23]. But this very maneuver undercuts the composite’s claimed value: if meaningful interpretation depends on dissecting the model to recover the original features, then the composite serves more as an obfuscation layer than a meaningful summary. This problem is especially acute when the model relies on abstract or minimally actionable features - such as principal components, DNA methylation sites, or gene expression levels. These inputs are often biologically opaque, and identifying them as “important” does little to clarify their clinical relevance. Even when the inputs include well-understood physiological measurements with known interventions, the case for using a composite remains weak. If actionable variables are already present, interpretation and clinical decision-making would be better served by focusing directly on those variables - without detouring through the opaque composite.

### 4.6 Emergent Composite Metrics

It is important to reiterate that the issue is not composite measures per se but the objectives they are used for in the “ageing” field. Like any model, their usefulness and validity are objective-dependent. The contradictions in how these measures have been applied become clearer when contrasted with “emergent” composite measures - those that arise naturally rather than being statistically constructed from a set of features. Such measures are common in nature, acting as low-cost heuristics that summarize a complex space of variables. One illustrative example is the concept of “fitness indicators” from evolutionary research. In sexual selection, certain traits serve as proxies for overall fitness. Mating requires choosing individuals capable of survival, reproduction, and adaptation to current pressures. The peacock’s tail exemplifies this: maintaining a large, symmetric, and vibrant tail is energetically expensive and demands multiple physiological systems to function well. Only peacocks that meet these demands can sustain such a display, making the tail an emergent measure of fitness that can be assessed at a glance. If any underlying function is compromised, the tail reflects it, offering a fast heuristic of “fitness”. Yet, crucially, while the tail summarizes fitness, it does not allow for inference of the precise combination of underlying traits or the history of events that gave rise to it. If a peacock’s tail deteriorates, it does not inherently indicate whether the cause is malnutrition, infection, or genetic defects - only that something has gone wrong. These natural “fitness indicators” highlight two key advantages of composite measures. First, they support decision-making without requiring a large set of variables to be directly measured - vital when data collection is costly or impractical. Second, even if every relevant trait could be measured, merging them into a meaningful composite poses a formidable challenge. Imagine if, rather than simply observing a peacock’s tail, a female had to measure each function - metabolic, immune, muscular, and more - and then combine them into a single “fitness score”. Not only would this be resource-intensive, but trying to predetermine the “best” integration of traits runs counter to the way evolution tests variation in ever-changing environments. Many common performance assessments can likewise be viewed as emergent composite measures capturing multiple systems’ integrated functions. Take a running test, for example: it depends on cardiovascular, respiratory, muscular, and neural coordination, effectively summarizing overall “function” without isolating each system’s role. If we tried measuring these systems individually - heart output, lung capacity, muscle strength, neural control, and so on - and then integrating them, we’d need a detailed grasp of the complex generative process that brings them together. That complexity can be prohibitively high, making holistic performance tests a more practical alternative^**17**^.

Emergent composite measures can also be useful as confounding proxies in statistical analyses [56]. In many research settings, understanding relationships among variables requires adjusting for others that are difficult or impossible to measure directly. One well-known example of such a confounding composite is chronological age. Researchers rely on age to capture a broad range of influences, especially when these cannot be specifically measured [3]. Although the “limitations” of chronological age spurred the development of better “biological age” composites, the difference in how they are employed is telling. Despite having access to the underlying features, they collapse them into a value and then promote it for various objectives. This contradiction is even more pronounced in predictive tasks: if you already have measures of each relevant “biological damage” marker and want to predict disease risk, does it make sense to first compress all that information into a single index only to then attempt to predict the outcome? For example, if muscle mass, visceral fat, and subcutaneous fat are measured, does it make sense to collapse them into total weight - or some weighted sum - and use that to predict outcomes? It depends on the objective [1, 32]. If the goal is to check an elevator’s weight limit, such aggregation is appropriate. But if the aim is to assess fat-related health risk, collapsing distinct variables can obscure critical information.

### 4.7 Composite Metrics and The Utility of Labels

Despite the fundamental mismatch between these composite measures’ properties and their intended objectives, their enduring and increasing popularity presents an interesting contradiction. This discrepancy merits careful examination, as it suggests either the limitations outlined so far are incorrect or these models serve purposes beyond their stated objectives. Based on my particular exposure to these models and the subset of literature I have examined, I find the latter explanation far more plausible^**18**^. What follows is necessarily a conjecture - an attempt to infer the alternative functions that might account for their persistence. In research, as in other domains, rhetoric and vision can sustain ideas well beyond what empirical scrutiny would justify, especially in biology, where complexity vastly outstrips understanding. Models built on vague yet evocative and simplistic concepts thrive because they create an illusion of coherence, offering a heuristic framework that seems to explain a wide range of observations while accommodating virtually any experimental result.

The endurance of these “ageing” composites, I suspect, is not a testament to their effectiveness in addressing their stated objectives, but rather to their rhetorical strength and interpretative flexibility. Perhaps the clearest illustration can be found in the so-called “second-generation clocks” - or, more precisely, in how they are typically presented and interpreted. In essence, these models are conventional time-to-event predictors of all-cause mortality. The novelty of these “technologies” is a simple scale transformation where the outcome is remapped onto an age scale: the score is expressed as “the chronological age at which an average individual would exhibit the same predicted risk” [23]. Contrasting the two representations of this same predicted quantity - raw risk versus age-transformed score - is informative. Although the only supported interpretation by either is a mortality risk, the age-transformed score enjoys additional advantages. The one commonly referenced in the literature is providing a “contextualized” and “easily comprehensible” estimate of mortality or disease risk [23]. At first glance, the claim seems plausible - everyone has an intuitive feel for age - but it does not withstand closer scrutiny. First, risk equivalence is not etiological or actionable equivalence. Two individuals may share the same predicted mortality risk, but the generative processes underlying that risk - and the clinical strategies required to address it - may be entirely distinct. A 25-year-old with end-stage renal failure and an “average 75-year-old” may both exhibit the same probability of death within a given time frame, yet expressing this as the younger individual having “the mortality of a 75-year-old” offers no meaningful clinical context [57]. It does not indicate a shared etiology, nor does it guide intervention in a way that the original risk estimate could not. Even among individuals who are chronologically 75, there is no single, coherent clinical profile to serve as a reference point (**Appendix I**). The notion of an average “75-year-old” context is, at best, a statistical abstraction - one that obscures the wide heterogeneity found in real-world populations. Referring to a younger patient as having “the mortality of a 75-year-old” invokes a context that is not meaningfully defined, offering the illusion of interpretability while failing to convey any concrete clinical insight. Second, the communicative benefit of converting risk into an “equivalent age” is overstated. Clinicians in routine care and clinical-trial settings already interpret hazard ratios, survival probabilities, absolute-risk differences and numbers-needed-to-treat; these metrics underpin day-to-day decisions in oncology, cardiology, critical care and beyond. Furthermore, clinicians do not assess risk in isolation; they interpret it within the broader clinical context, which already includes the patient’s age along with other variables such as comorbidities.

When pressed, the developers of these scales often insist that the purpose of the age transformation is purely communicative - to represent mortality risk in a more intuitive form and not to suggest any deeper biological equivalence (personal communication). Yet the way these models are routinely presented, both in academic literature and broader discourse, suggests otherwise. Far from being treated as a mere statistical convenience, the transformed scores are often framed as explanatory entities: proxies for “systemic ageing processes”, measures of “biological age”, or even candidate “ageing biomarkers”. Visualizations, language choices, and comparative framing all contribute to this shift. Age-equivalent scores are displayed in plots that label deviations as “accelerated” or “decelerated ageing” [58, 59], while papers and press releases routinely tout findings such as “participants were biologically 4 years younger”. In this way, a scale transformation becomes a narrative engine, animating claims far beyond what the underlying model justifies. This rhetorical elasticity [60, 61, 62, 63, 64] delivers tangible dividends - it enhances the model’s visibility, marketability, and funding appeal. Unlike a straightforward prediction of mortality, the label “biological age” suggests broader relevance across diverse health domains, allowing the metric to be linked to virtually any condition. After all, if “ageing” is framed as “damage accumulation” [10], then any condition with some form of “deterioration” will be relevant. This conceptual promiscuity makes it adaptable to various disease areas, policy initiatives, and funding opportunities. Crucially, all these advantages come at little cost as the construct these scores claim to represent is inherently latent and vaguely defined. This ambiguity shields it from falsification^**19**^. If two second-generation clocks yield diverging results, it is said they reflect different “dimensions of ageing”. If they agree, it is taken as mutual validation. If an intervention alters one of these scores, it is framed as evidence of its effectiveness; if it does not, the result is a clue about unmeasured pathways. In every case, the framing ensures its rhetorical utility regardless of outcome. This ensures that these labels remain beyond empirical contestation, residing purely in the realm of rhetoric and definition. After all, the model is designed not just to measure the construct, but to define it.

## Acknowledgements

This publication is based on publicly available research data from The Baltimore Longitudinal Study of Aging study (BLSA) that has been made available through the Alzheimer’s Disease Data Initiative Workbench. The BLSA is supported by the Intramural Research Program (IRP) of the National Institute on Aging (NIA). The authors thank the investigators, staff and participants of the BLSA study for making the data available. BLSA investigators have not contributed to nor approved, and are not in any way responsible for, the contents of this publication. The author thanks the Alzheimer’s Disease Data Initiative for access to the AD Workbench used in this study.

## 5. Appendix I - On Average Ageing

This article originated as a brief note on the concept of “average ageing”, inspired by an older piece titled “The Average Man” [65]. That article discussed the 1950 U.S. Air Force Anthropometric Survey, which sought to design equipment and cockpits suitable for the broad range of Air Force personnel by measuring around 11 physical dimensions. The survey found that out of thousands of pilots, virtually none fit within the average range across all measured dimensions. This high-lighted the challenges of employing average concept for multivariate design. This finding was interesting as a similar principle underlies many constructs in the “ageing” field - particularly those aimed at quantifying “biological age” or “age deviation”. For example, as defined in a recent perspective article: “Conceptually, an individual’s age defined by the level of age-dependent biological changes, such as molecular and cellular damage accumulation. In practical use, this is often summarized as a number (in units of time) matching the chronological age where the average person in a reference population shares the individual’s level of age-dependent biological changes” [10]. Given the variability and weak correlations often observed among ageing-related changes [66, 57, 67, 68, 8, 10, 69], and the large number of features typically used to construct composite metrics, the likelihood of any individual being “average” across all variables is exceedingly low. This is reflected in assumptions (A4) Common Base-line and (A5) Shared Trajectory, which underlie many ageing models. Initially, this article set out to examine the plausibility of these assumptions. However, it quickly became clear that this was the wrong question [32]. Models should be assessed based on their utility over a set of given objectives, rather than whether they ‘exist’ in a literal sense. After all, the notion of “average” can be operationalized in many ways. One could define a sample as average if it falls within a fixed range around the mean of all variables simultaneously, or if it meets that criterion for a subset (e.g., 80%) of variables. The acceptable range itself could also vary by feature. Moreover, “average” can be treated not categorically (average vs. non-average), but dimensionally - quantifying how average a profile is [1]. Each of these definitions constitutes a model, and like any model, its validity must be judged in light of a specific and clearly defined objective [32]. This is why A4 and A5 received less focus in this article. Instead, attention was directed toward whether the latent construct of “biological age” adds meaningful utility for various objectives, and toward more foundational assumptions - (A1) through (A3) - that critically shape that utility.

- **(A4) Common Baseline**: These models implicitly assume that individuals begin life at or near a common baseline value of the latent construct.
- **(A5) Shared Trajectory**: They also assume that the latent construct progresses along a common normative trajectory across individuals.

As with other assumptions in latent-variable frameworks, A4 and A5 are not directly testable, since the latent construct is itself unobservable. Consequently, debates around their validity often rest on rhetorical appeals to plausibility rather than empirical adjudication. More critically, even if A4 and A5 were true, deviation-based composites would remain subject to the same structural limitations discussed throughout this work - especially feature convergence (A1). For that reason, our analysis centers on the shared constraints (A1–A3), with A4 and A5 noted here for completeness but not examined in depth.

These categories serve as convenient labels for discussion but do not represent truly distinct domains. As with any categorization, these labels impose boundaries on a multidimensional space and are defined based on what is of interest.

**Figure S1.**
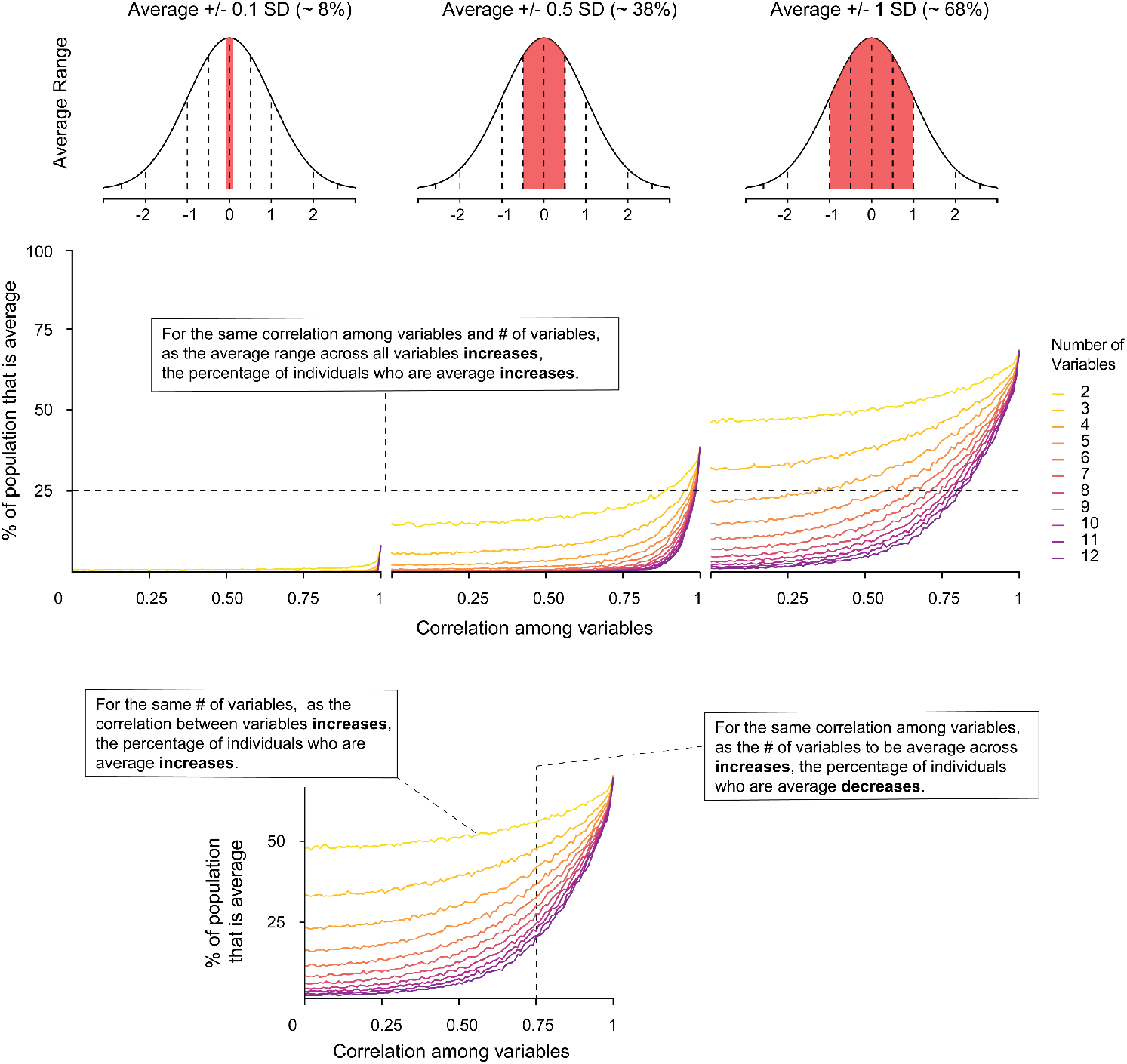
Simulation illustrating how the proportion of samples meeting the criteria for the “average” label changes with adjustments to the range around the mean, number of variables, and correlations among variables. As the average range broadens, a higher percentage of samples are deemed “average”. Conversely, for a given average range, increasing the number of variables and/or decreasing variable correlations reduces the likelihood of a sample being classified as “average” across all dimensions. Samples were drawn from multivariate normal distributions using the function ‘mvrnorm’ from the R package MASS. The number of samples was set to 10000. The means of all variables was set to 0. The covariance matrix was specified with variances set to 1 along the diagonal, and correlations (off-diagonal elements) ranging from 0 to 1, with a step size of 0.01. A sample was considered ‘average’ if the value of each variable fell within a specified range around the mean, defined as mean ± range * sd.

**Figure S2.**
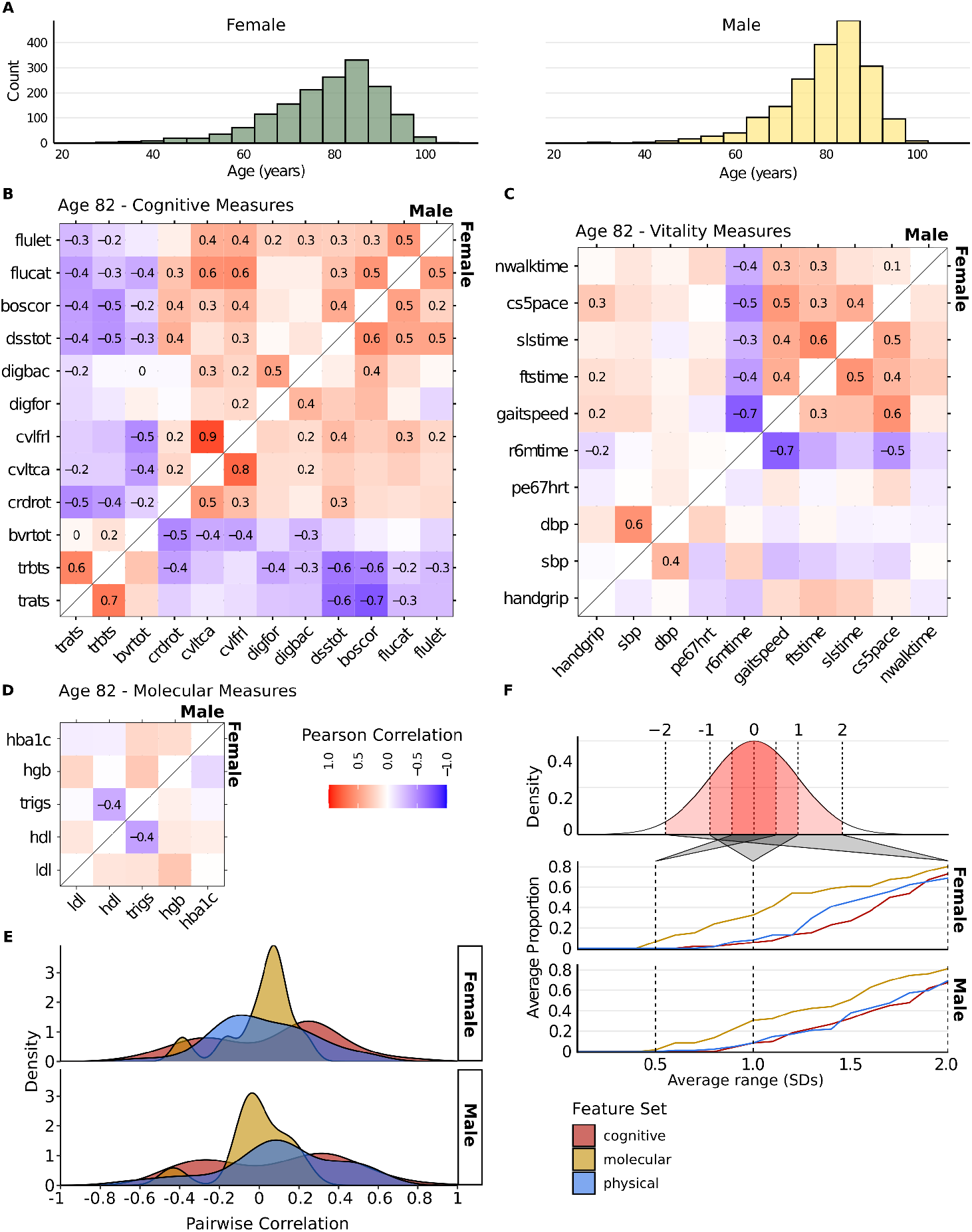
Characterization of an 82-year-old participant from the BLSA Open Data. **(A)** Histogram of age of female (left) and male (right) participants. **(B)** Heatmap of Pearson correlations among selected subset of cognitive measures at age 82, separated by sex. **(C)** Heatmap of Pearson correlations among selected subset of vitality measures at age 82, separated by sex. **(D)** Heatmap of Pearson correlations among selected subset of molecular measures at age 82, separated by sex. **(E)** Distribution of pairwise-correlations at age 82 stratified by sex. The x-axis represents pairwise correlation coefficients, while the y-axis shows the density of these correlations. Different colors indicate distributions of correlations for the three considered sets of features. **(F)** Proportion of individuals classified as “average” (standardized values for all measures falling between the mean ± the specified range) for ranges spanning 0 to 2 standard deviations, shown separately for females (middle panel) and males (lower panel). The Benjamini-Hochberg procedure for multiple testing correction was applied separately by sex and within each set of measures (cognitive, vitality, and molecular). Correlation values are displayed as text only for associations with an adjusted p-value < 0.05.

**Figure S3.**
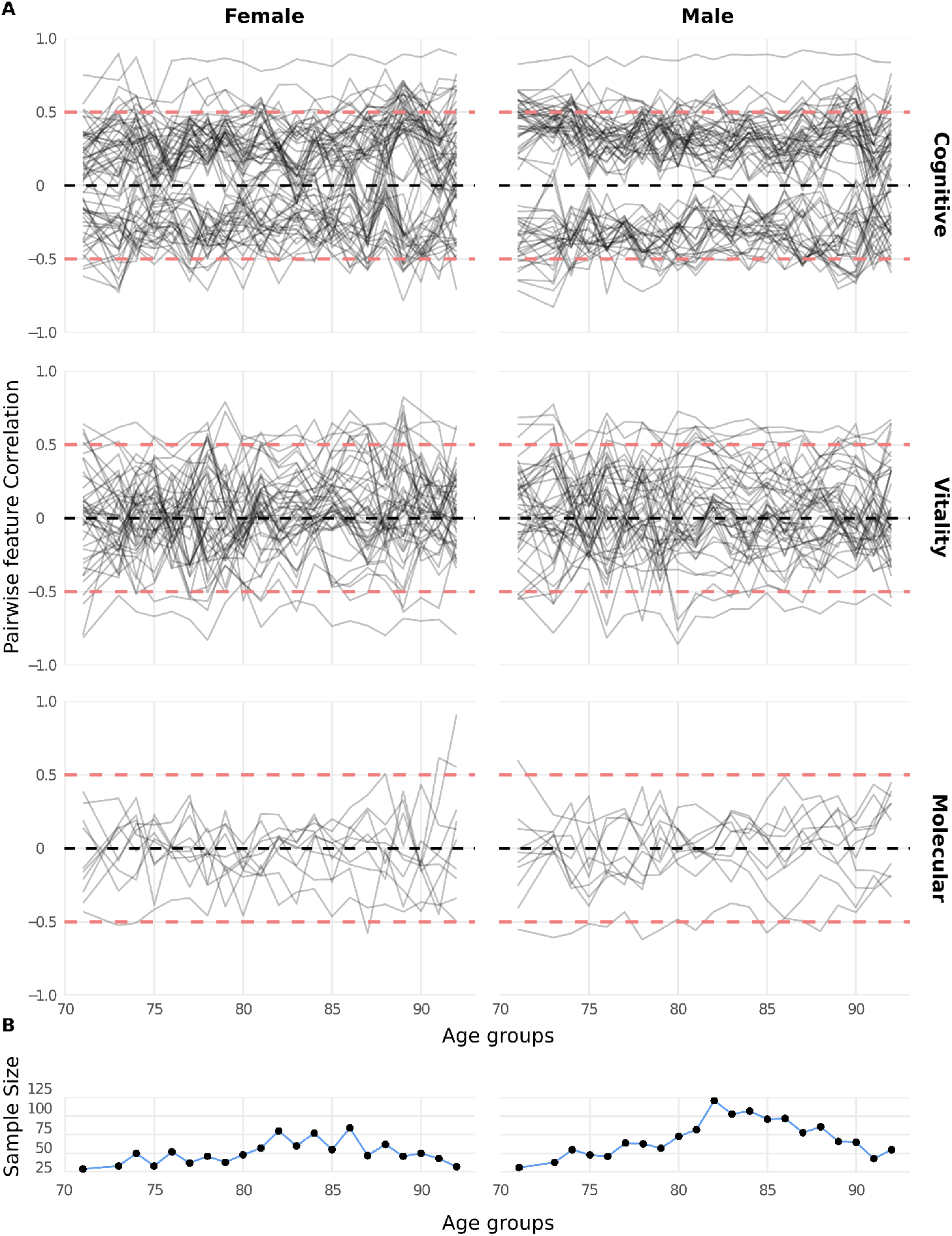
**(A)** Pairwise feature correlations across three measurement sets (cognitive, vitality, and molecular) for different age groups (71–92 years), stratified by sex (female on the left, male on the right). Each line represents the correlation for a specific feature pair across age groups. The y-axis shows the correlation values ranging from -1 to 1. The black dashed line indicates a correlation of 0, while the red dashed lines represent a correlation of +/-0.5. **(B)** Sample sizes for each age group, stratified by sex (female on the left, male on the right). The y-axis indicates the number of individuals in each age group, and the x-axis corresponds to the age groups (71–92 years).

**Table S1.**
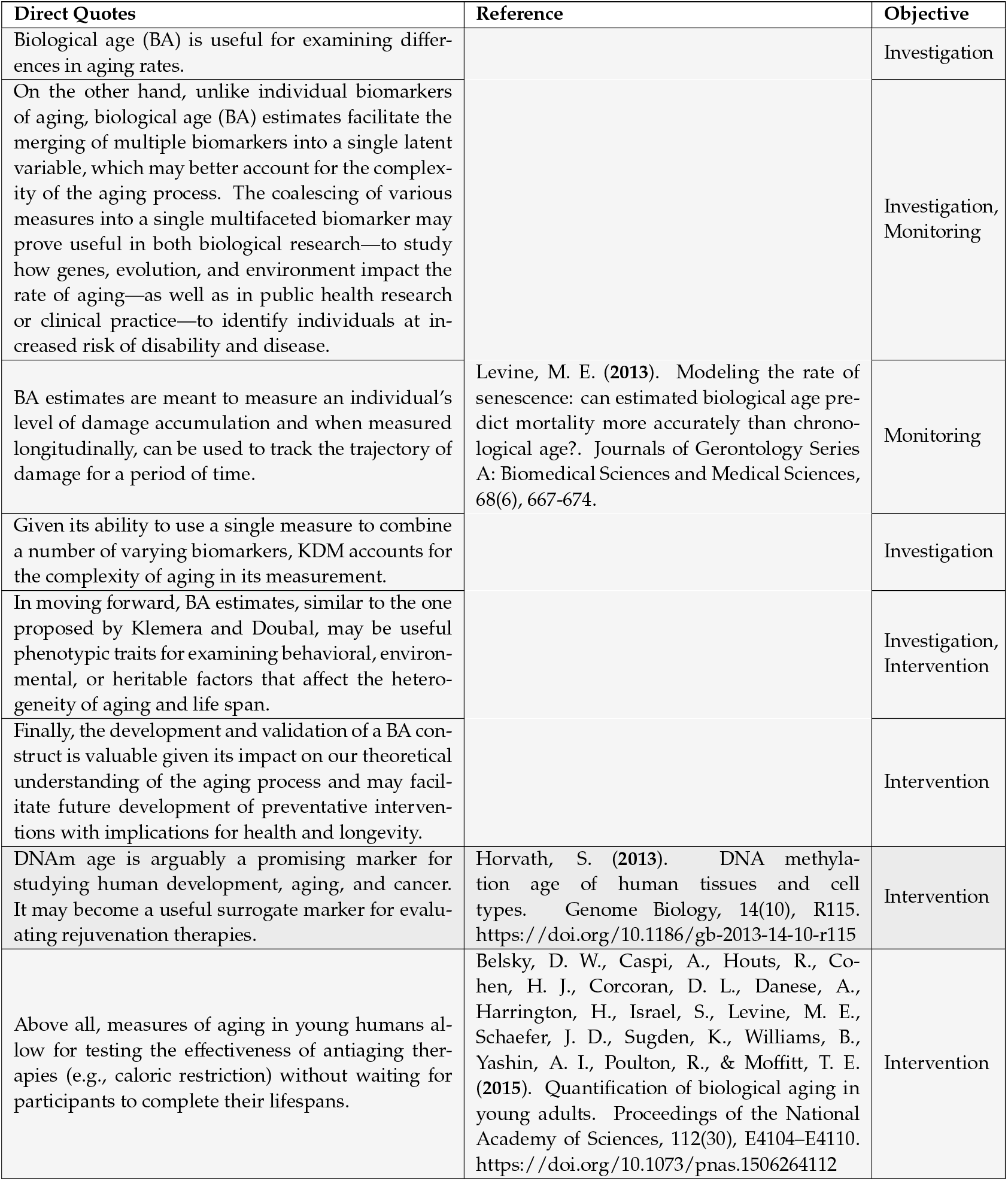
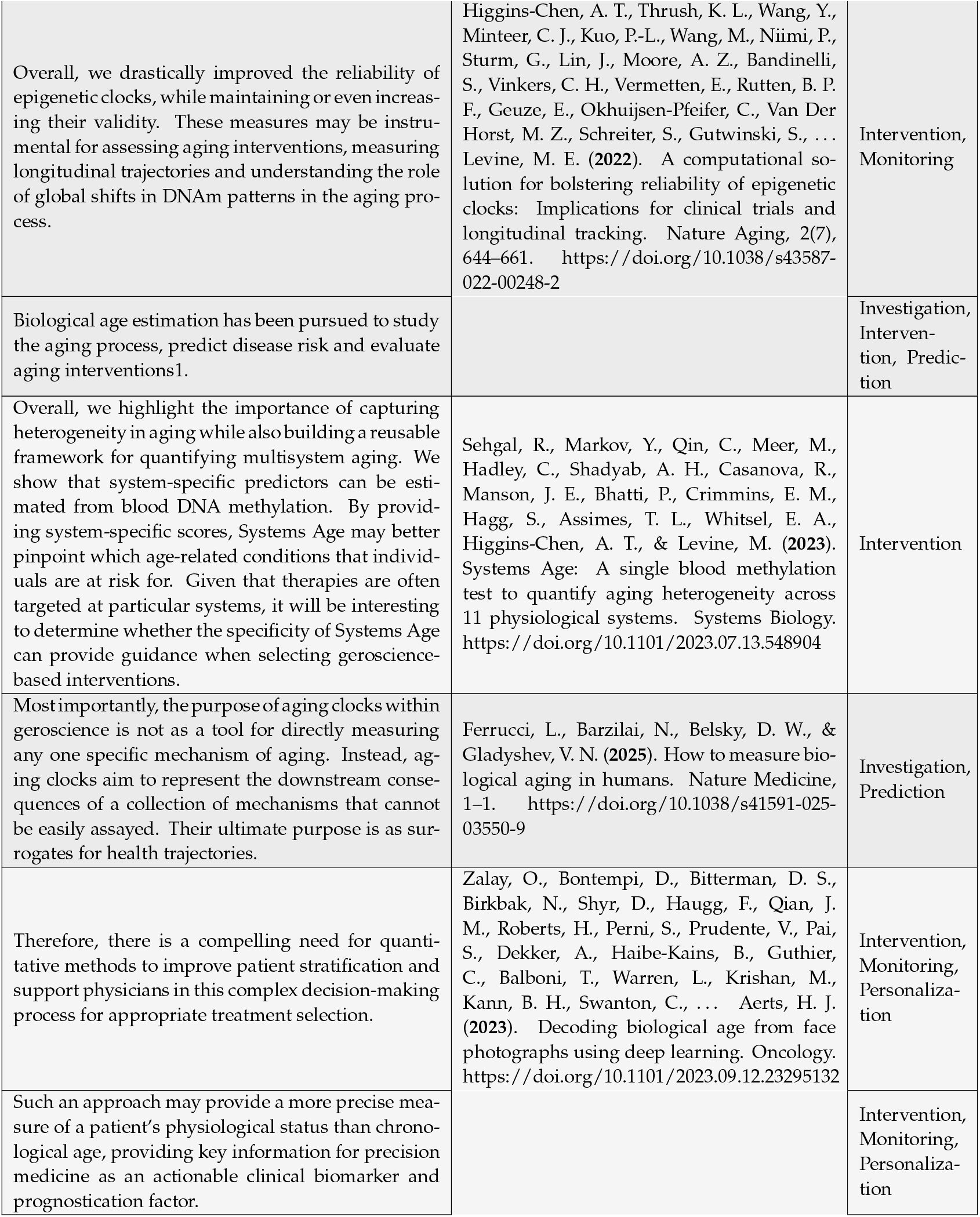
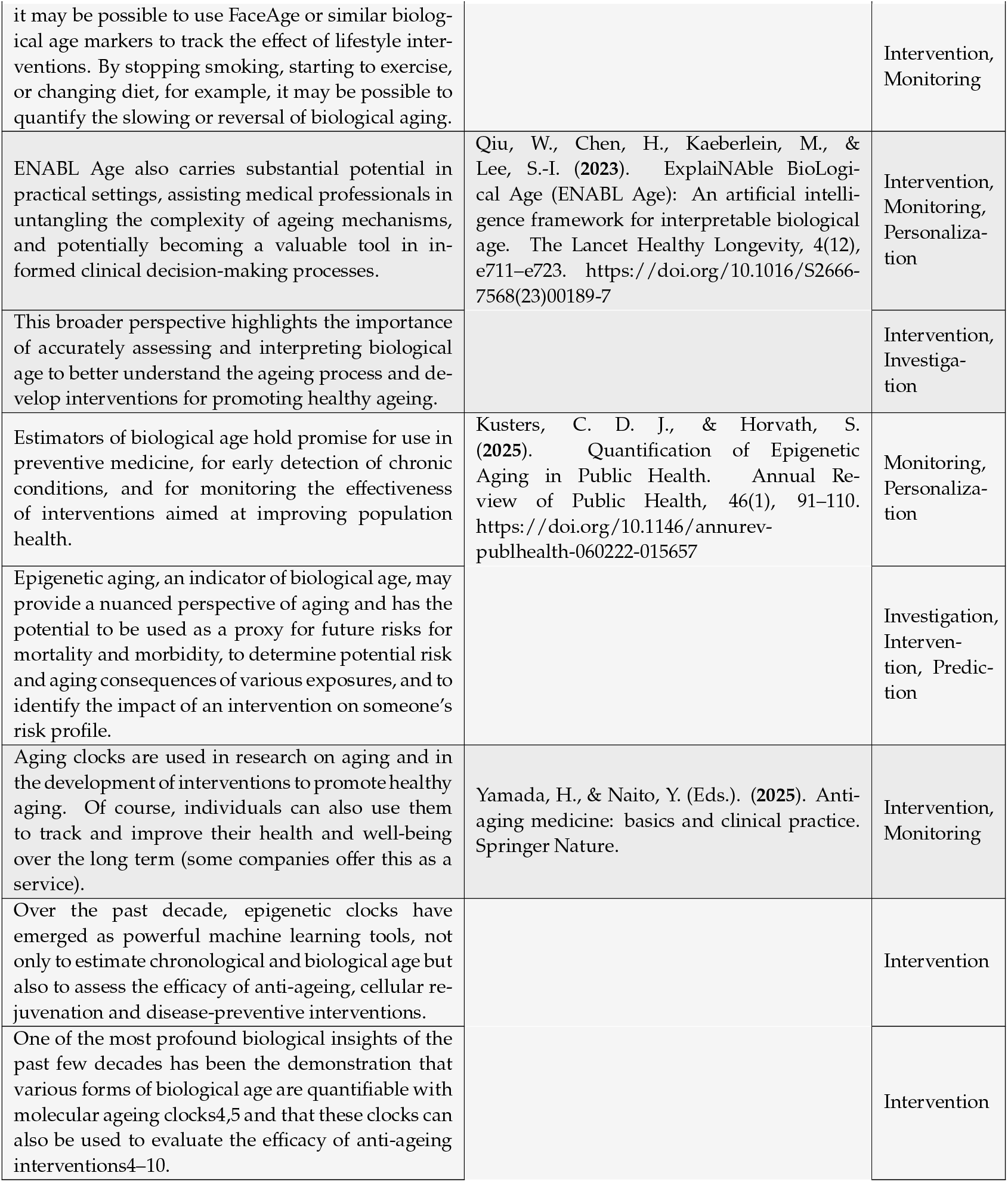
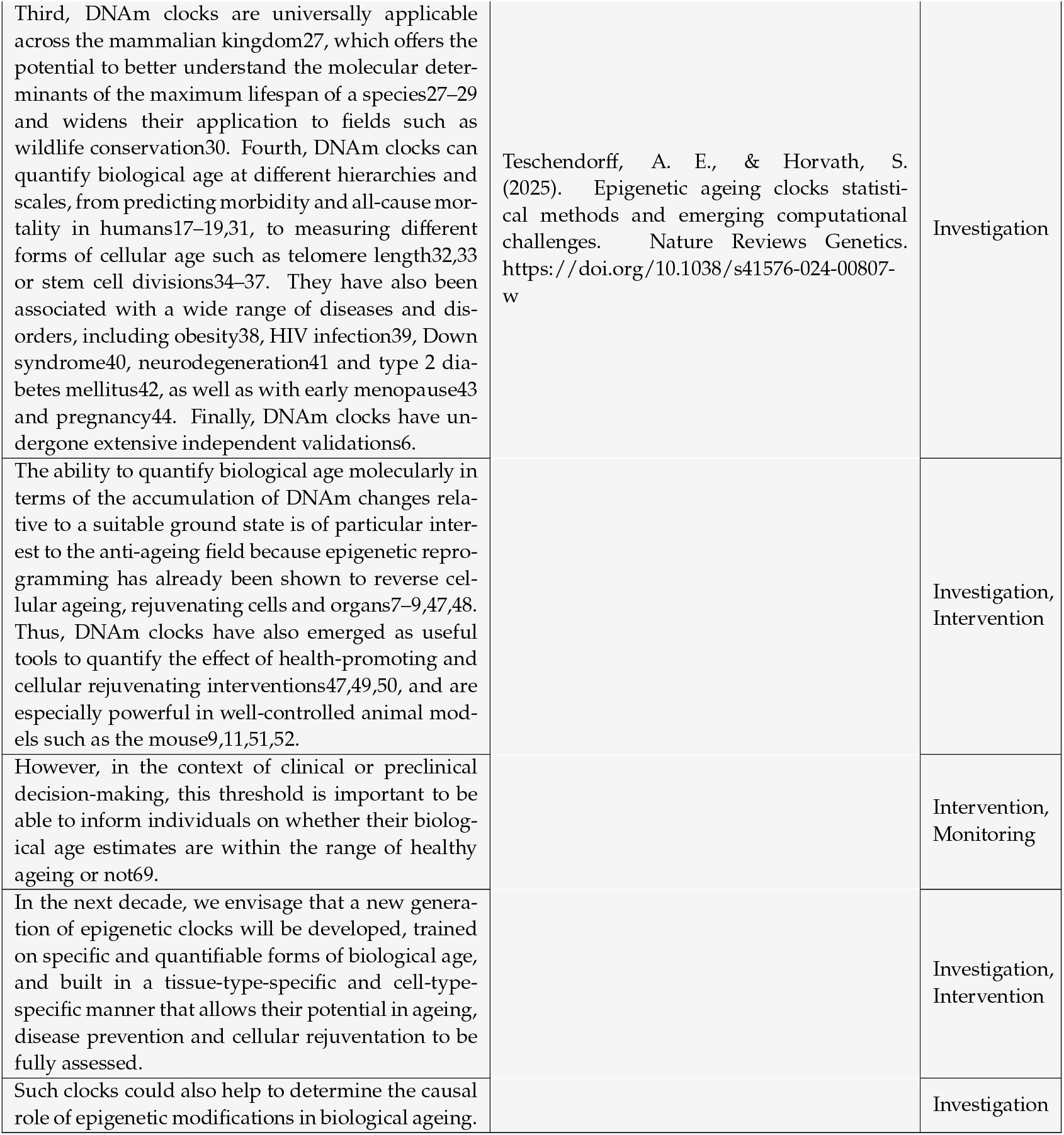
Illustrative quotes highlighting the objectives for which composite “ageing” metrics are advocated.

**Table S2.**
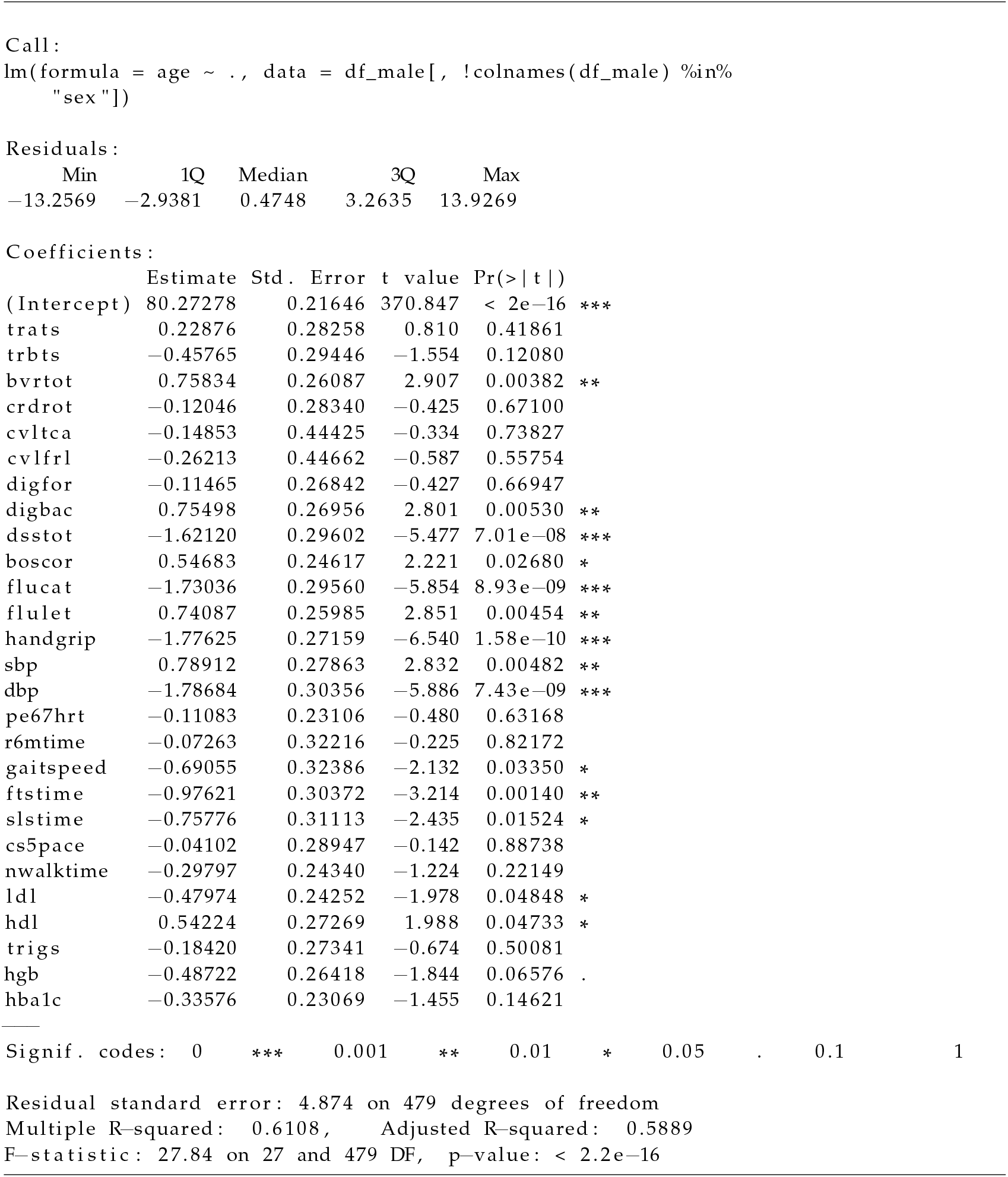
Summary of linear model predicting age from 27 features in males from BLSA Open Data.

**Table S3.**
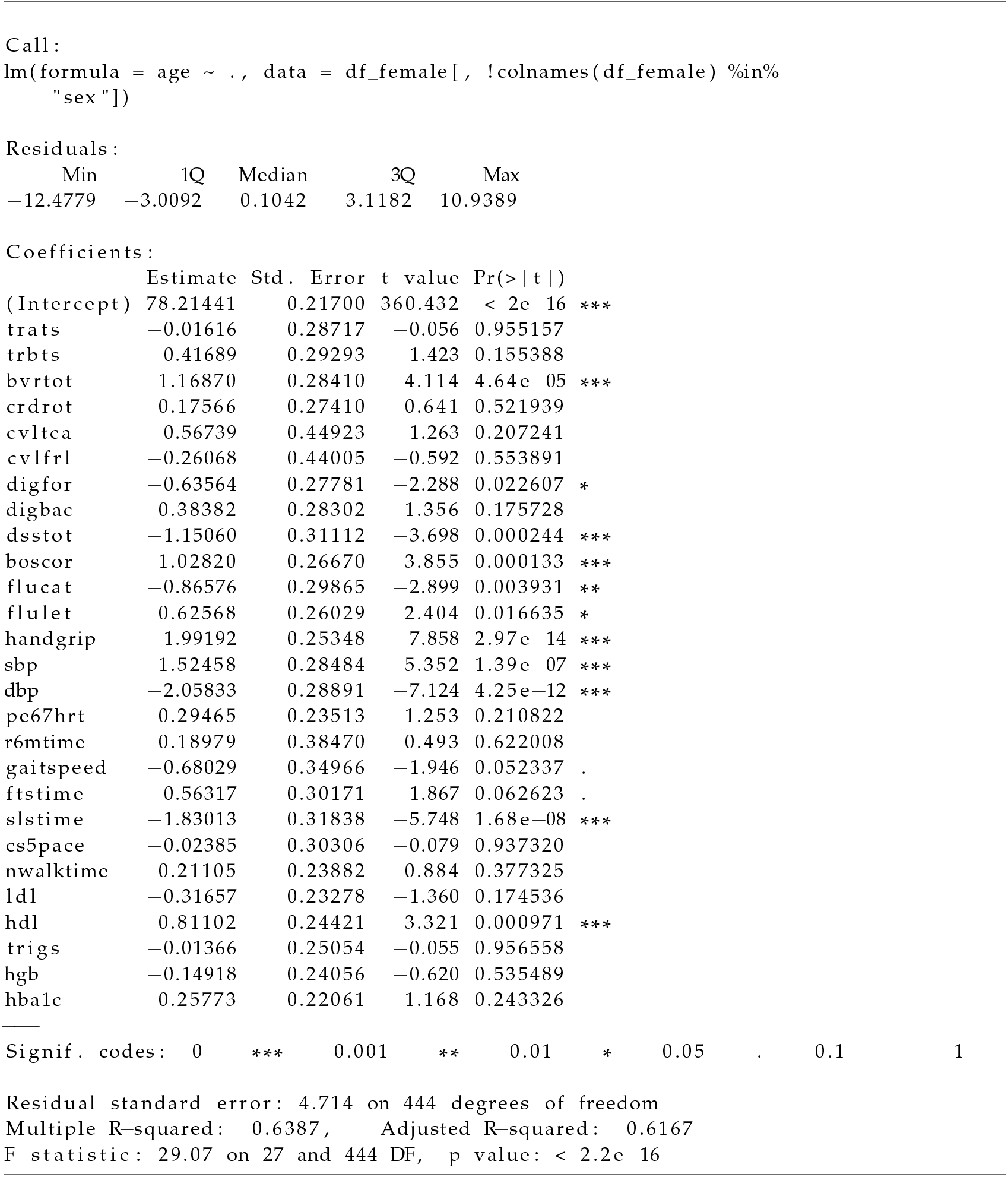
Summary of linear model predicting age from 27 features in females from BLSA Open Data.

## Footnotes

1 **Terminology**: Throughout this article, composite variable refers to the empirical score derived from observable features, while latent construct refers to the unobservable property—such as “biological age” or “health state”—that the composite is intended to represent.

2 The selection of input features (predictors) and distal outcomes (endpoints) is inherently contentious within latent-construct frameworks. Because the constructs themselves are unobservable, debates about what qualifies as a valid input or outcome often rely on rhetorical justification—appeals to biological plausibility, clinical relevance, or theoretical coherence—rather than empirical necessity. While these discussions shape how a model is framed, they do not alter the core assumptions or structural properties of the generative model examined here. The issues explored in this article—such as feature convergence, loss of information, and the limits of latent-variable inference—persist regardless of which specific features or outcomes are chosen. For that reason, these definitional debates are acknowledged but not addressed in detail.

3 The degree of convergence is contingent on the covariance structure of the input features. Some might argue that this issue can be “mitigated” by selecting perfectly correlated features or designing a model that yields a unique outcome for every possible input combination. However, such strategies contradict the assumptions of the latent construct model - and undercut the very rationale for using composite scores in the first place. A fuller treatment of this point is provided in the Discussion section.

4 **Terminology**: Throughout this article, “theoretical model” refers to the posited generative process in which a latent construct gives rise to both proximal and distal measures. “Statistical model” refers to an empirical implementation of this structure using observed data (“Statistical Model A”) that operationalizes the relationships among variables.

5 The outcome(s) used to train a statistical model are not always conceptualized as distal outcomes in the generative sense (i.e., events or states that unfold downstream of the latent construct). Instead, they are often chosen simply because they are believed to be informative about the latent construct itself. For example, chronological age is frequently used as a training target in models of “biological age”, not because it is a distal effect of biological aging, but because it is assumed to correlate with the latent construct. This distinction is important: the statistical model does not require the outcome to be causally downstream, only that it serves as a useful proxy. However, the validity of this assumption strongly influences how the resulting composite is interpreted and applied.

6 Models that operationalize age deviation - the difference between an individual’s predicted criterion value and the expected value for their chronological age - not only inherit the structural limitations of the base latent-construct framework (A1–A3), but also introduce additional, often implicit, assumptions:

7 These categories serve as convenient labels for discussion but do not represent truly distinct domains. As with any categorization, these labels impose boundaries on a multidimensional space and are defined based on what is of interest.

8 This property appears under various labels across fields: degeneracy, equifinality, basins of attraction, non-identifiability and observable equivalence. Despite the differing terms, they all capture the same core behavior: multiple distinct configurations producing indistinguishable outputs.

9 Molecular-based “ageing” composites gained popularity for seemingly enabling more direct cross-species comparisons [36]. Yet, this apparent advantage is often illusory, as differences in how molecular features combine to produce function across species undermine direct comparisons.

10 For example, visceral fat (fat surrounding internal organs) or fat deposited within muscles can carry different health risks than subcutaneous fat (fat under the skin).

11 In practice, “Ageing” metrics do more than serve as proxies - they often end up defining the very concept of “ageing” they are meant to measure. This circularity reinforces the vagueness of the target construct and further complicates interpretation.

12 This problem is particularly pronounced in BA composites built from molecular features, which are often non-intuitive and difficult to interpret on their own. As a result, tracing what a change in the composite actually reflects becomes even more challenging.

13 Heterogeneity can be found at nearly every level of biological organization—from organs and tissues to cells and molecular profiles. Simply moving to finer resolutions does not guarantee more meaningful or actionable insight. As measurement granularity increases, so too does the difficulty of separating signal from noise. More importantly, organisms are structured hierarchically: low-level variation is integrated, buffered, or transformed through complex regulatory architectures and dynamical constraints. These include phenomena akin to “basins of attraction” in dynamical systems, where different microstates converge toward similar macro-level outcomes. In such settings, what appears as heterogene-ity at one level may be irrelevant or even misleading for a given objective at another. Thus, the resolution at which variation is measured—and the relevance of that variation—should be determined in relation to the specific goal of the analysis, rather than assumed to improve with increased detail.

14 In practice, a broader endpoint can outperform a narrowly defined one when the specific outcome is recorded inconsistently or is inherently noisy. For instance, overall-survival data are usually more reliable than progression-free-survival data;

15 The irony is that deep learning models are often introduced with reference to their ability to capture non-linear and high-dimensional relationships. Yet rather than using this capacity to improve prediction of meaningful outcomes, they are instead deployed to condense features into a one or a set of “age” composites.

16 Latent construct models also afford considerable flexibility [15] and many degrees of freedom for interpretation and post-hoc theory retrofitting. This interpretive flexibility will be examined in detail in a later section.

17 The difficulty of integrating disparate variables grows the farther one moves “down” the biological hierarchy. This challenge is particularly amplified for molecular features (e.g., genetic variants, DNA methylation patterns, or gene expression levels). Each intervening layer introduces additional nonlinear interactions and context-specific effects, compounding uncertainty and making a principled aggregation of measures ever more difficult [50, 51, 52, 53, 54, 55].

18 This view also aligns with the recurring research trends discussed earlier in this article, which are difficult to account for unless the structural limitations - specifically in applying these models to their promoted objectives - are indeed valid.

19 This vagueness also facilitates the repeated rebranding of the underlying concept. Over time, the same core idea has been repackaged under fresh labels - “functional age”, “biological age”, “ageing clocks”, “frailty index”, “biomarkers of ageing”. Each repackaging presents the illusion of progress, allows for renewed attention and for rhetorical distancing from accumulating observations of their limitations. By shifting terminology, proponents can refresh interest in the same underlying approach while avoiding direct engagement with prior criticisms

